# Myofibroblast Ccn3 is regulated by Yap and Wwtr1 and contributes to adverse cardiac outcomes

**DOI:** 10.1101/2022.12.01.518714

**Authors:** Michael A. Flinn, Santiago Alvarez-Argote, Makenna C. Knas, Victor Alencar Almeida, Samantha J. Paddock, Xiaoxu Zhou, Tyler Buddell, Ayana Jamal, Pengyuan Liu, Jenny Drnevich, Michaela Patterson, Brian A. Link, Caitlin C. O’Meara

## Abstract

While Yap and Wwtr1 regulate resident cardiac fibroblast to myofibroblast differentiation following cardiac injury, their role specifically in activated myofibroblasts remains unexplored. Here we assess the pathophysiological and cellular consequence of genetic depletion of Yap alone (*Yap^fl/fl^;Postn^MCM^*) or Yap and Wwtr1 (*Yap^fl/fl^;Wwtr1^fl/+^;Postn^MCM^*) in adult mouse myofibroblasts following myocardial infarction and identify and validate novel downstream factors specifically in cardiac myofibroblasts that mediate pathological remodeling. Following myocardial infarction, depletion of Yap in myofibroblasts had minimal effect on heart function while depletion of Yap/Wwtr1 resulted in smaller scars, reduced interstitial fibrosis, and improved ejection fraction and fractional shortening. Single cell RNA sequencing of interstitial cardiac cells 7 days post infarction showed suppression of pro-fibrotic genes in fibroblasts derived from *Yap^fl/fl^*,*Wwtr1^fl/+^;Postn^MCM^* hearts. In vivo myofibroblast depletion of Yap/Wwtr1 as well in vitro knockdown of Yap/Wwtr1 dramatically decreased RNA and protein expression of the matricellular factor Ccn3. Administration of recombinant CCN3 to adult mice following myocardial infarction remarkably aggravated cardiac function and scarring. CCN3 administration drove myocardial gene expression of pro-fibrotic genes in infarcted left ventricles implicating CCN3 as a novel driver of cardiac fibrotic processes following myocardial infarction.

## Introduction

Following cardiac injury such as myocardial infarction (MI), progressive fibrosis can contribute to adverse ventricular remodeling and ultimately heart failure. Resident cardiac fibroblasts, which directly produce the pro-fibrotic response, are a heterogeneous population of stromal cells. When activated following injury, a subset of resident fibroblasts adopt a myofibroblast phenotype (1, 2) characterized by expression of Periostin (Postn) and alpha smooth muscle actin (αSMA (1)), increased production of extracellular matrix (ECM) proteins (primarily collagens (3, 4)), and secretion of metalloproteases (1), inflammatory chemokines, and cytokines (5, 6). While myofibroblast function is critical for wound contraction, initial scar formation, and to prevent catastrophic heart rupture (1, 7), extended myofibroblast proliferation and activation can lead to persistent matrix protein secretion causing exacerbated scar formation, myocyte uncoupling, decreased cardiac compliance, and progressive heart failure (5). There is currently an un-met need to identify molecular mediators of myofibroblast activity so that we can therapeutically target these pathways and regulate the fibrotic response following cardiac injury.

The Hippo-Yap/Wwtr1 pathway (herein referred to as ‘Hippo-Yap’) is a highly conserved signaling pathway consisting of core protein kinases; Stk3, Stk4, Lats1, and Lats2, and scaffolding proteins; Sav1, Mob1a, and Mob1b, whose kinase activity suppresses function of the transcriptional co-activators Yap and Wwtr1 (8). Phosphorylation of Yap and Wwtr1 by the Hippo pathway kinases prevents nuclear localization and promotes phospho-degradation of Yap and Wwtr1, thereby inhibiting nuclear transcriptional activity. It has been demonstrated in mice Lats1/2 deletion specifically in resident cardiac fibroblasts promotes Yap and Wwtr1 activity, driving proliferation and cell fate transition to a myofibroblast state in a cell autonomous manner and resulting in deleterious fibrosis in mice (11, 12). Depletion of Yap/Wwtr1 in Tcf21 or Collagen (Col1a2 or Col11a1) positive cells attenuated pathological remodeling and improved cardiac function (11–14). However, Tcf21, Col1a2 and Col11a1 target all fibroblast populations and the role of endogenous Yap and Wwtr1 in differentiated cardiac myofibroblasts specifically has not been investigated. Here, we take the novel approach of investigating the role of endogenous Yap/Wwtr1 expression in activated cardiac myofibroblasts (Postn+) following MI. We demonstrate a cooperative role for myofibroblast Yap and Wwtr1 in scar size, interstitial fibrosis, and cardiac function post MI.

The downstream pathways modulated by Hippo-Yap activity in myofibroblasts are poorly understood. We found that depletion of Yap/Wwtr1 in myofibroblasts both in vivo and in vitro strongly suppresses expression of the secreted matricellular protein Ccn3 (also known as nephroblastoma overexpressed— Nov), suggesting Ccn3 expression might contribute to adverse outcomes post MI. Indeed, global administration of recombinant CCN3 to mice post MI significantly exacerbates adverse cardiac function and scarring. CCN3 signaling to myocardial tissue post MI promoted expression of pro-fibrotic genes. Collectively, we identify a novel factor expressed downstream of Yap/Wwtr1 in cardiac fibroblasts that independently promotes adverse cardiac wound healing dynamics.

## Results

### Yap and Wwtr1 are expressed in myofibroblasts after MI

After an ischemic injury, resident cardiac fibroblasts differentiate into myofibroblasts which aggregate at the site of injury as soon as 1 dpi with a robust presence around 4-7 dpi (1). By comparison, uninjured hearts show little signs of myofibroblast differentiation (<1% of interstitial cells). To assess whether the Hippo-Yap pathway was active in myofibroblasts, we leveraged a myofibroblast specific and tamoxifen inducible Postn Cre (*Postn^MCM^* (1)) crossed to a transgenic reporter line containing a floxed stop codon prior an enhanced green fluorescent protein (eGFP) in the Rosa26 locus (*R26-eGFP^fl/+^*) to express eGFP in myofibroblasts following tamoxifen induction (**Figure 1A**). This reporter model facilitates reliable identification of myofibroblasts by eGFP expression in tissue sections. *R26-eGFP^fl/+^*;*Postn^MCM^* mice were crossed to Yap and Wwtr1 floxed mice to generate Yap^fl/fl^;Wwtr1^fl/wt^;*R26-eGFP^fl/+^*; *Postn^MCM^* reporter mice that are depleted for Yap and Wwtr1 expression in Postn expressing cells following tamoxifen administration (**Figure 1B**). We performed MI followed by tamoxifen treatment in both postnatal day 6 (P6) and in 8-10 week old adult mice to investigate expression patterns of Yap and Wwtr1 at two developmental timepoints (**Figures 1C,D**). At 7 dpi we identified a robust population of GFP+ cells at the site of injury, but not in remote cardiac tissue, in *R26-eGFP^fl/+^*;*Postn^MCM^* mice (**Figures 1E,G, and S1**). At both timepoints we observed nuclear Yap and Wwtr1 expression in GFP+ myofibroblasts in *R26-eGFP^fl/+^*;*Postn^MCM^* mice while *Yap*^fl/fl^;*Wwtr1*^fl/+^;*R26^eGF^*;*Postn^MCM^* mice showed marked decrease in nuclear Yap (**Figures 1E,F**) and Wwtr1 (**Figures 1G,H**) expression.

**Figure 1.**
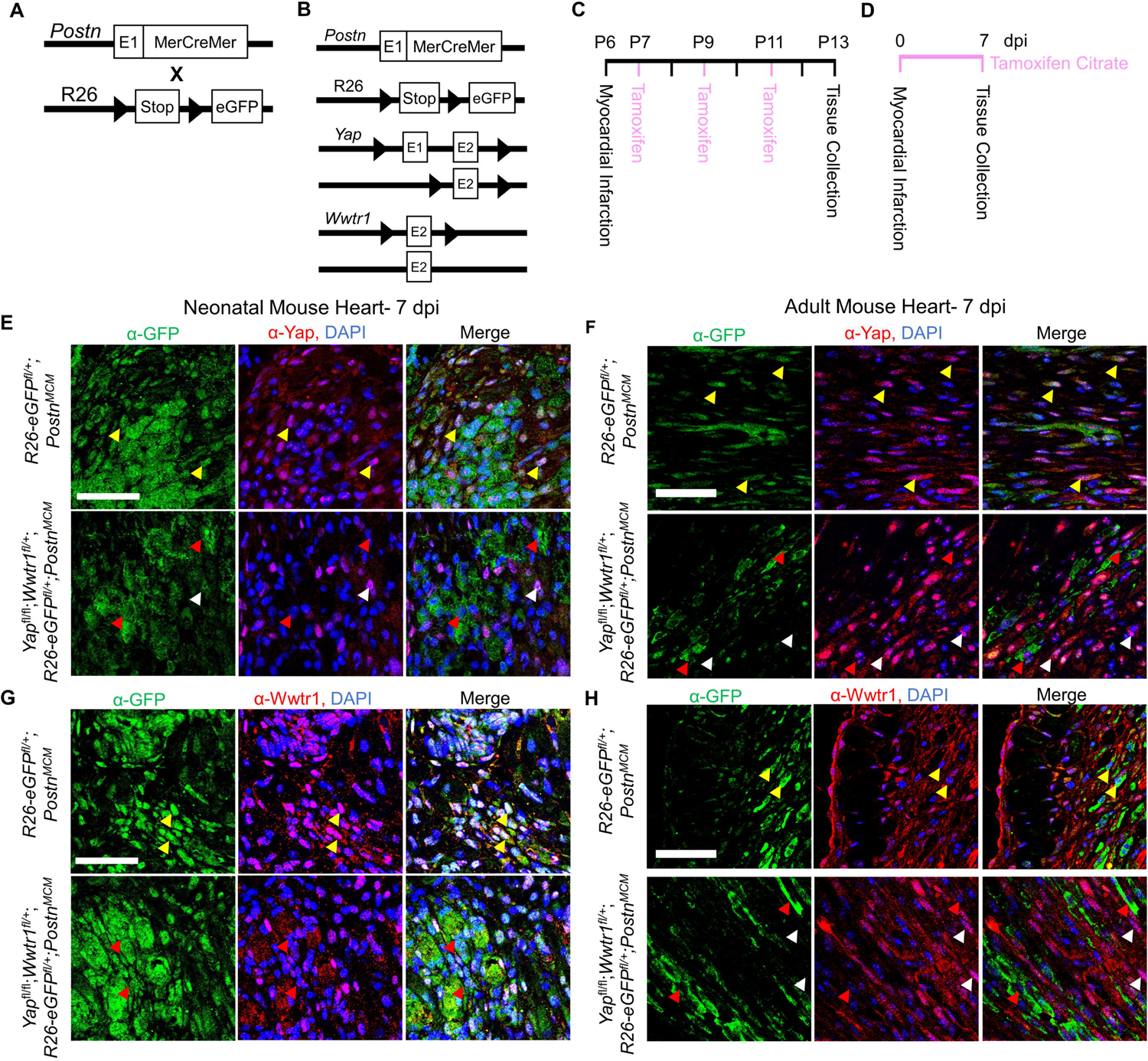
Yap and Wwtr1 expression is detected in myofibroblasts after ischemic injury. (A) Schematic representation of the alleles present in *R26-eGFP^fl/+^;Postn^MCM^* and (B) *Yap^fl/fl^;Wwtr1^fl/+^;eGFP^fl/+^;Postn^MCM^* animals. (C) Experimental timeline of P6 or (D) adult MIs and subsequent tamoxifen administration. (E through H) Representative confocal images of the scar region in *R26-eGFP^f/+^;Postn^MCM^* and *Yap^fl/fl^;Wwtr1^fl/+^;R26-eGFP^f/+^*;*Postn^MCM^* mice at 7 dpi. Scale = 100 µm. (E and G) denote P13 juvenile mice while (F and H) are adults. (E and F) Immunostaining with α-GFP and α-Yap antibodies and DAPI. Yellow arrows denote Yap co-localization in GFP positive cells, red arrows denote absence of Yap colocalization in GFP positive cells, white arrows denote nuclear localized Yap in GFP negative cells. (G and H) Immunostaining with α-GFP and α-Wwtr1 antibodies and DAPI. Yellow arrows denote Wwtr1 co-localization in GFP positive cells, red arrows denote absence of Wwtr1 colocalization in GFP positive cells, white arrows denote nuclear localized Wwtr1 in GFP negative cells.

### Depletion of *Yap* in myofibroblasts results in modest protection against ventricular dilation post MI

To test if Yap deletion in myofibroblasts influences cardiac function we performed MI on *Yap^fl/fl^;Postn^MCM^* and *Yap^fl/fl^* animals followed by tamoxifen administration provided *ad libitum* in chow. We found no difference between genotypes in cardiac function at baseline (3 days before MI) or at 3 dpi indicating similar degree of injury in both groups (**Figure S2A**). At 28 dpi *Yap^fl/fl^;Postn^MCM^* animals had significantly smaller left ventricular internal diameters during diastole (LVID-d) compared to *Yap^fl/fl^* (**Figures S2A,B**). LVID-s by comparison was not significantly different (p = 0.27). While the *Yap^fl/fl^;Postn^MCM^* genotype conferred modest protection against left ventricular dilation, %EF and %FS were not significantly different between *Yap^fl/fl^;Postn^MCM^* and *Yap^fl/fl^* animals (**Figure S2A**) nor were heart weights or body weights (**Figures S2C,D**). At the histological level we found no significant difference in CM cross sectional area between experimental and control animals, suggesting reduced ventricular dilation does not correlate with attenuated CM hypertrophy in our model (**Figures S2E,F**). Together, these data illustrate that Yap depletion from myofibroblasts imparts a slight protection against adverse ventricular dilation after MI, but functional contractility parameters such as %EF and %FS remain comparable to control animals.

Considering *Yap^fl/fl^;Postn^MCM^* animals showed a slight improvement in ventricular dilation following injury, we sought to investigate if cell proliferation or scarring differences could be observed. To assess cell cycle activity, following MI we administered a single dose of EdU at 6 dpi, a time when *Postn* expression and myofibroblast proliferation is high in the heart (1, 2). We found a 70% decrease in EdU+ nuclei in the scar region of *Yap*^fl/fl^*;Postn^MCM^* mice compared controls (**Figures S2G,H**). Based on the known role of Yap in promoting cell proliferation, we postulated that the significant reduction of scar associated cell EdU incorporation is the result of decreased myofibroblast proliferation. However, despite the reduction in purported myofibroblast proliferation, we did not observe any change in scar size after injury as assessed by both midline scar percentage and the percentage of collagen content within the left ventricle of Gömöri trichrome stained serial sections (**Figures S2I,J**).

### Depletion of both Yap and Wwtr1 in myofibroblasts improves cardiac function following MI

Yap and Wwtr1 share overlapping roles in various cell types. In some cases, depletion of Yap can be compensated by increased Wwtr1 activity and vice versa (15). In vitro knockdown of either Yap or Wwtr1 in neonatal rat cardiac fibroblasts demonstrated a significant decrease in DNA synthesis compared to negative siRNA or non-treated cells while an additive effect was observed with knockdown of both genes suggesting redundant role for Yap and Wwtr1 in fibroblasts (**Figure S3**). We hypothesized that depletion of both Yap and Wwtr1 in myofibroblasts would improve cardiac outcomes post MI compared to Yap depletion alone. To test this hypothesis, we used a genetic mouse model were both copies of *Yap* and a single copy of *Wwtr1* are floxed, as both *Postn^MCM^* and *Wwtr1* are located chromosome 3 in mice at approximately 3 million base pairs apart. This genetic linkage made generating a homozygous floxed *Wwtr1* line with the Cre driver locus impractical, either alone or in combination with *Yap*.

We subjected *Yap*^fl/fl^;*Wwtr1*^fl/+^;*Postn^MCM^* or *Yap*^fl/fl^;*Wwtr1*^fl/+^ littermate controls to MI and subsequently fed a continuous diet of tamoxifen citrate chow (**Figure 2A**). Similar to the loss of Yap alone, depletion of Yap and Wwtr1 did not affect heart weight or body weight at 60 dpi (**Figures 2B,C**). While we found no difference in cardiac function between genotypes at baseline or 3 dpi, *Yap*^fl/fl^;*Wwtr1*^fl/+^;*Postn^MCM^* mice showed significantly improved cardiac function at 60 dpi compared to control (**Figures 2D,E**). While %FS and %EF declined to ∼10% and ∼17% respectively in *Yap*^fl/fl^;*Wwtr1*^fl/+^ animals, %EF and %FS in *Yap*^fl/fl^;*Wwtr1*^fl/+^;*Postn^MCM^* were maintained at ∼18% and ∼35% respectively. Both LVID-s and LVID-d also trended downward in *Yap^fl/fl^;Wwtr1^fl/+^;Postn^MCM^* mice at 60 dpi, but were not statistically different from control mice (p = 0.17 and 0.22 respectively, **Figure S4**). In contrast, when allowed to progress to the 60 dpi timepoint, *Yap^fl/fl^*;*Postn^MCM^* still showed no difference in %EF or %FS or fibrosis compared to *Yap^fl/fl^*littermates and heart weight and body weight were not affected (**Figure S5,S6,S7**). We tested if the preservation in cardiac function in *Yap^fl/fl^;Wwtr1^fl/+^;Postn^MCM^* as compared to *Yap^fl/fl^;Wwtr1^fl/+^* controls was the result of Cre expression in within myofibroblasts alone by repeating the MI study on *Postn^MCM^* transgenic mice and wildtype littermate controls. We found no difference in %FS or %EF between these genotypes indicating that preserved cardiac function is due to deletion of Yap and Wwtr1, and not due to Cre expression in myofibroblasts (**Figure 2E**). Together, these data illustrate that depletion of both Yap and Wwtr1 attenuate adverse cardiac remodeling and improve cardiac function after ischemic injury.

**Figure 2.**
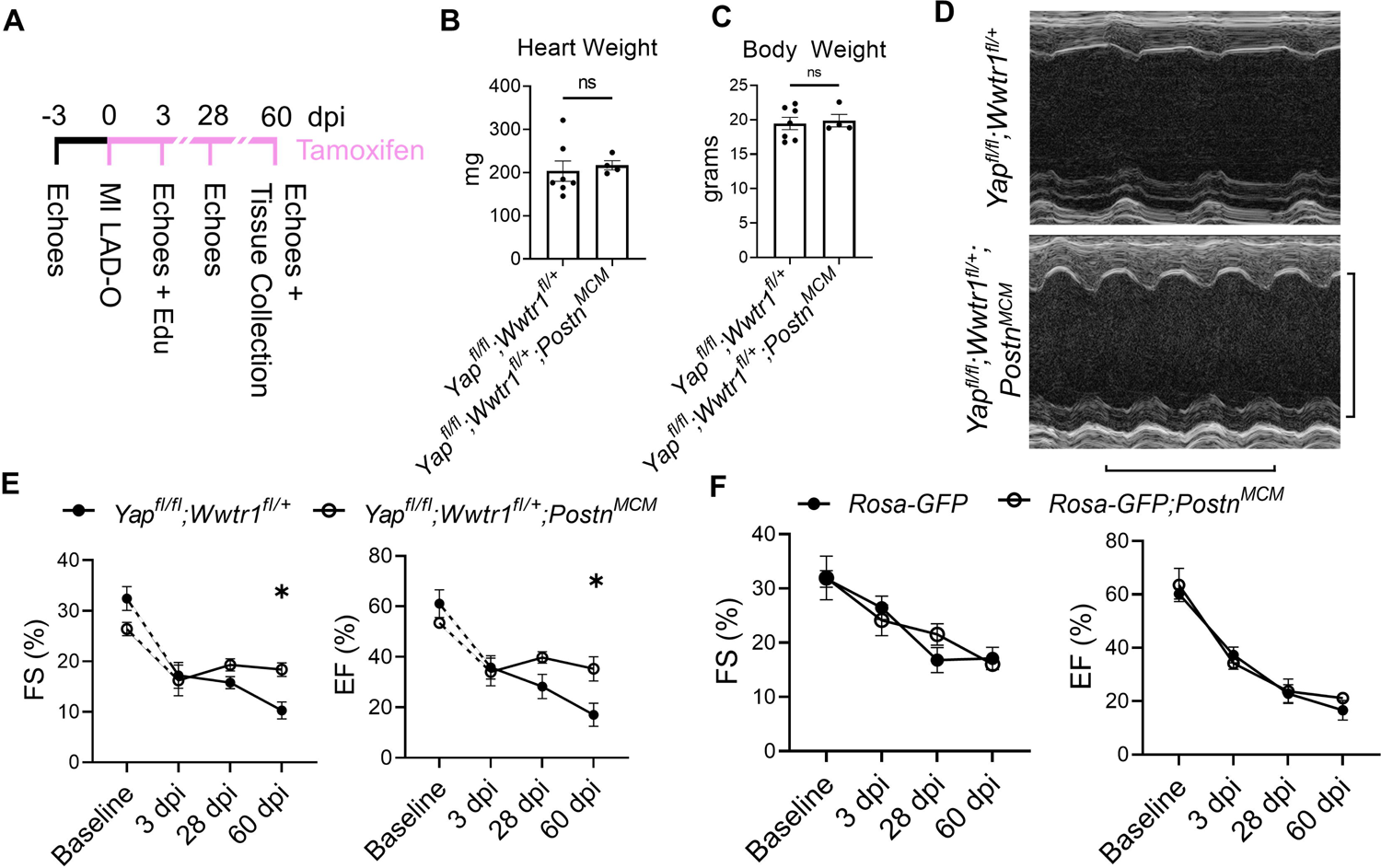
Depletion of Yap and a single copy of Wwtr1 from myofibroblasts improves left ventricular function in response to ischemic injury. (A) Experimental timeline of echocardiography, MIs, tamoxifen chow administration, and EdU administration in adult *Yap^fl/fl^;Wwtr1^fl/+^* and *Yap^fl/fl^;Wwtr1^fl/+^;Postn^MCM^* mice. Quantification of (B) heart and (C) body weights at 60 dpi. n = 7 *Yap^fl/fl^;Wwtr1^fl/+^* and 4 *Yap^fl/fl^;Wwtr1^fl/+^;Postn^MCM^*. Unpaired student’s t-test. (D) Representative M-mode echocardiograms of left ventricles 60 dpi. Short axis view at mid papillary muscle. Horizontal bar = 500 ms. Vertical bar = 5 mm. (E and F) Quantification of %FS and %EF analyzed by repeated measures two-way ANOVA and Sidak multiple comparisons test. n = 7 *Yap^fl/fl^;Wwtr1^fl/+^*, 4 *Yap^fl/fl^;Wwtr1^fl/+^;Postn^MCM^*, 5 *R26-eGFP^f/+^*, 4 *R26-eGFP^f/+^;Postn^MCM^*. A separate subset of animals establishing a baseline for *Yap^fl/fl^;Wwtr1^fl/+^*animals as denoted by the hashed lines and assessed by un-paired student’s t-test. n = 4 *Yap^fl/fl^;Wwtr1^fl/+^* and 5 *Yap^fl/fl^;Wwtr1^fl/+^;Postn^MCM^*. ns = not significant, * = P <0.05.

Despite the improved cardiac function of *Yap^fl/fl^;Wwtr1^fl/+^;Postn^MCM^* mice after surgical MI, no difference in cardiomyocyte cross sectional area was observed between genotypes (**Figures 3A,B**). We next tested whether depletion of both Yap and Wwtr1 from myofibroblasts resulted in modulation of cell proliferation, scar formation, or fibrosis in our model. To quantify proliferation of cells within the scar region, adult mice subjected to MI were administered a single dose of EdU at 3 dpi. Consistent with results observed in *Yap^fl/fl^;Postn^MCM^* mice, *Yap*^fl/fl^;*Wwtr1*^fl/+^;*Postn^MCM^* mice exhibited a 60% decrease in EdU+ scar associated cells compared to controls, whereas the percentage of EdU+ interstitial cells remained unchanged (**Figures 3C,D and S8**). However, unlike *Yap^fl/fl^;Postn^MCM^* mice which showed no difference in scar size at 28 or 60 dpi (**Figures S2I,J and S5,S6**), *Yap^fl/fl^;Wwtr1^fl/+^;Postn^MCM^* mice displayed significantly reduced scar size as assessed by scar midline length (38% reduction) and fibrotic percentage of the left ventricle (38% reduction) (**Figures 3E,F**) and a 43% reduction in interstitial fibrosis compared to control (**Figure 3G,H**). We further characterized scar composition by quantifying denatured collagen, which is more easily turned over and can reduce deleterious fibrosis in the heart, using collagen hybridizing peptide Cy3 conjugate (CHP) (16). CHP binds to the unfolded triple-helix of collagen fibers, thus marking denatured collagen. CHP has been shown to correlate with other assays assessing degradation of collagen post MI such as matrix metalloproteinase activity in vivo and zymography (17, 18). While scars from *Yap*^fl/fl^*;Postn^MCM^* mice showed no difference in CHP binding compared to controls (**Figures 3I,J**)*, Yap*^fl/fl^;*Wwtr1*^fl/+^;*Postn^MCM^* mice displayed ∼2.5 fold increase in the amount of denatured collagen in the scar region as compared to *Yap*^fl/fl^;*Wwtr1*^fl/+^ littermates (**Figures 3K,L**). Thus, depletion of both Yap and Wwtr1 significantly modulated scar size and also collagen composition following MI.

**Figure 3.**
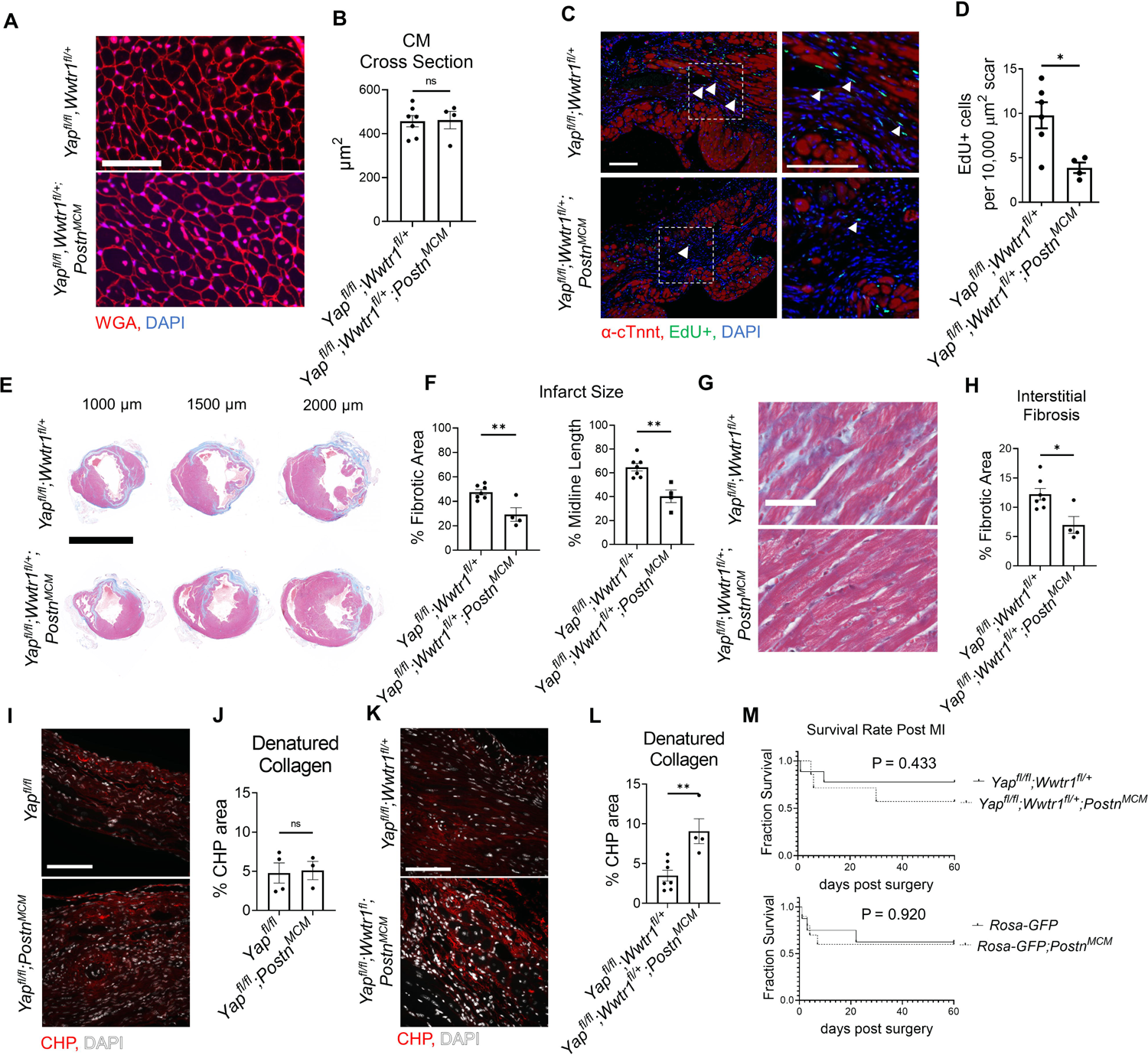
Depletion of myofibroblast Yap and Wwtr1 decreases fibrosis at 60 dpi. (A) Representative images of WGA staining in the remote zone of the left ventricle at 60 dpi. Scale = 50 µm. (B) Quantification of cardiomyocyte cross sectional area. n = 7 *Yap^fl/fl^;Wwtr1^fl/+^* and 4 *Yap^fl/fl^;Wwtr1^fl/+^;Postn^MCM^*. Unpaired student’s t-test. (C) Representative images denoting EdU+ nuclei within the scar area. White arrows indicate EdU+ scar associated nuclei. Scale = 100 µm. (D) Quantification of EdU+ scar associated nuclei. n = 6 *Yap^fl/fl^;Wwtr1^fl/+^* and 4 *Yap^fl/fl^;Wwtr1^fl/+^;Postn^MCM^*. Unpaired student’s t-test. (E) Representative serial sections of Gömöri trichrome stained hearts measured from the apex. Scale = 5mm. (F) Quantification of infarct scar size by either total fibrotic area or midline size of the left ventricle. n = 7 *Yap^fl/fl^;Wwtr1^fl/+^* and 4 *Yap^fl/fl^;Wwtr1^fl/+^;Postn^MCM^*. Unpaired student’s t-test. (G) Representative images depicting interstitial fibrosis within the remote zone of Gömöri trichrome stained hearts. Scale = 50µm. (H) Quantification of the % blue fibrotic area versus total ventricular tissue. n = 7 *Yap^fl/fl^;Wwtr1^fl/+^* and 4 *Yap^fl/fl^;Wwtr1^fl/+^;Postn^MCM^*. Unpaired student’s t-test. (I,K) Representative images of CHP staining within the scar region of 60 dpi mice. Scale = 100 µm. (J,L) Quantification of CHP as a % area of the scar region. n = 4 *Yap^fl/fl^*, 3 *Yap^fl/fl^;Postn^MCM^*, 7 *Yap^fl/fl^;Wwtr1^fl/+^*, and 4 *Yap^fl/fl^;Wwtr1^fl/+^;Postn^MCM^*. Unpaired student’s t-test. (M) Survival curve comparing *Yap^fl/fl^;Wwtr1^fl/+^* and *Yap^fl/fl^;Wwtr1^fl/+^;Postn^MCM^* or *R26-eGFP^f/+^* and *R26-eGFP^f/+^*;*Postn^MCM^* mice after MI until the 60 dpi end point. Animals that died during surgery were not included. n = 9 *Yap^fl/fl^;Wwtr1^fl/+^*, 7 *Yap^fl/fl^;Wwtr1^fl/+^;Postn^MCM^,* 8 *R26-eGFP^f/+^*, 10 *R26-eGFP^f/+^;Postn^MCM^.* Logrank Mantel-Cox test. ns = not significant, * = P <0.05, and ** = P <0.01.

Prior studies have demonstrated that genetic ablation of *Postn* expressing cells (i.e. myofibroblasts) results in a stark decreased survival post MI due to lack of scar deposition and subsequent rupture (1). Importantly, neither depletion of Yap and Wwtr1 nor expression of Cre itself significantly affect survival of infarcted mice (**Figure 3M**). Collectively, combined depletion of Yap and Wwtr1 in myofibroblasts attenuates fibrosis and improves functional outcomes following MI.

### Single cell analysis defines Yap and Wwtr1 downstream targets in cardiac myofibroblasts post MI

We next performed transcriptomic profiling on interstitial cells from *Postn^MCM^* and *Yap*^fl/fl^;*Wwtr1*^fl/+^;*Postn^MCM^* hearts post injury to identify transcriptional changes and differential infiltration of cell types between genotypes. At 7 days post MI, hearts were extracted, digested into single cell suspension, and FACS sorted for live nucleated cells. We targeted cDNA and library construction of 3,000 cells per genotype on the 10x Chromium Controller for sequencing. We obtained a total of 6,063 high quality sequenced cells, 2,930 from *Postn^MCM^* and 3,133 from *Yap*^fl/fl^;*Wwtr1*^fl/+^;*Postn^MCM^* hearts, at an average sequencing depth of 47,945 reads per cell. Cluster analysis of all cells revealed 18 main populations consisting of neutrophils (clusters 1, 18), B cells (cluster 2), macrophages/monocytes (clusters 0, 3, 11, 12, 13), T cells (clusters 4, 7, 8, 10), fibroblasts (cluster 5), natural killer cells (cluster 14), and dendritic cells (cluster 17) (**Figures 4A,B**, **Table S4**). Two small clusters (15 and 16) could not be identified or expressed mixed cell type markers. While cells from each genotype were represented in most clusters, clusters 6, 9, and 10 were primarily derived from *Yap*^fl/fl^;*Wwtr1*^fl/+^;*Postn^MCM^* (**Figures 4C**). Cluster 6 and 10 were enriched for proliferation markers (*Top2a* and *Mki67*), suggesting a population of proliferative T cells infiltrate myocardial injury in response to Yap/Wwtr1 myofibroblasts deletion.

**Figure 4.**
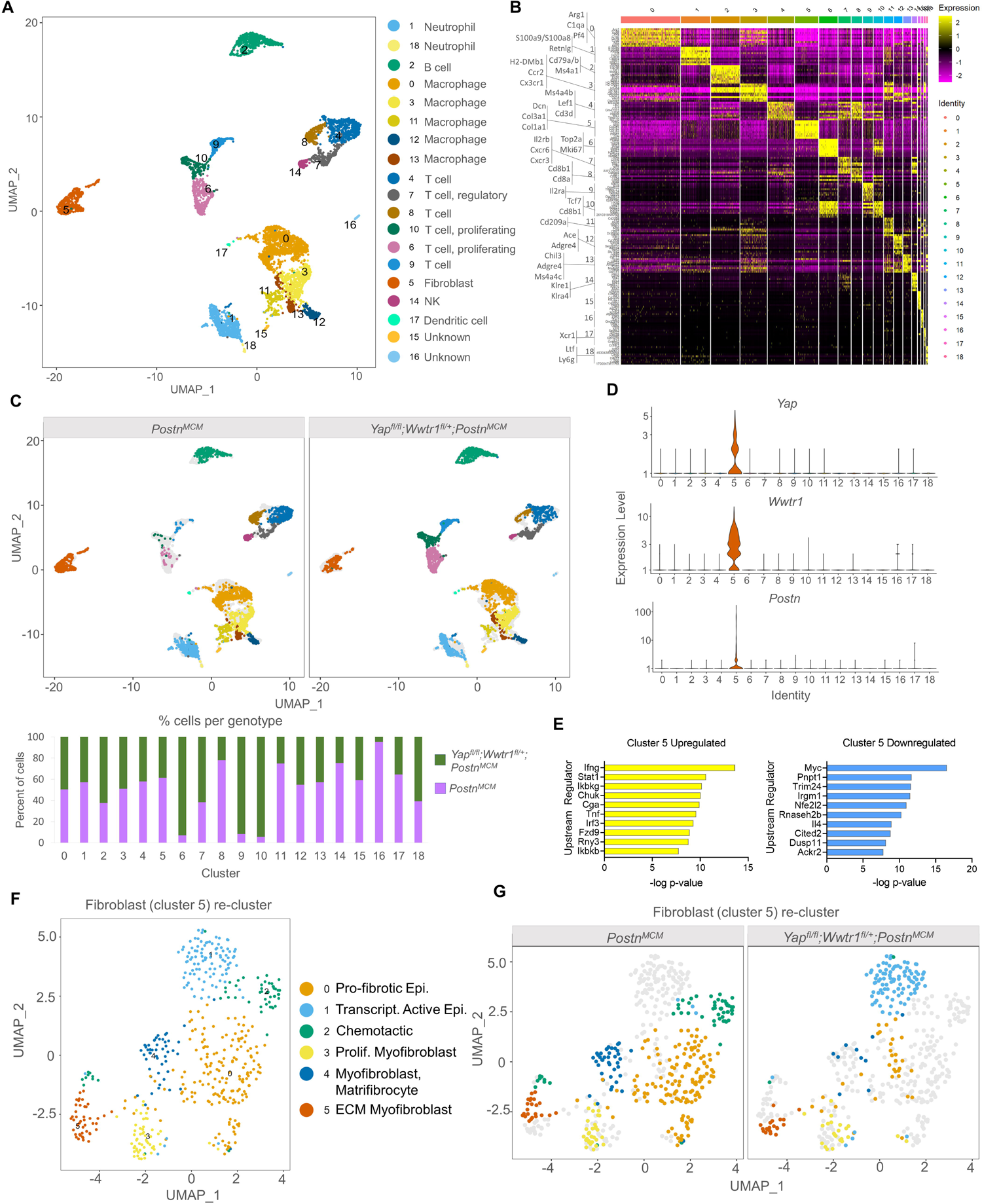
scRNAseq reveals Yap and Wwtr1 downstream targets in cardiac fibroblasts post MI. (A) UMAP projection and identification of combined interstitial cardiac cells, post filtering, from *Postn^MCM^* and *Yap^fl/fl^;Wwtr1^fl/+^;Postn^MCM^* adult mouse heart 7 dpi. Clusters are identified by enriched gene markers in Table S4. (B) Heatmap of top 10 genes enriched within each cluster. (C) UMAP projection split by genotype and quantification of percent of cells from each genotype for each cell cluster. (D) Violin plots denoting UMI count for *Yap*, *Wwtr1*, or *Postn* for each cluster. (E) IPA analysis of predicted upstream regulators of upregulated or downregulated DEGs (Y*ap^fl/fl^;Wwtr1^fl/+^;Postn^MCM^* versus *Postn^MCM^*) in fibroblasts (cluster 5). (F) UMAP projection of re-clustering only fibroblasts (cluster 5) and identification of cardiac fibroblast subclusters. Epi.= epicardial, Trascript. = transcriptionally, Prolif. = Proliferating, Myofib. = myofibroblast, MFC= matrifibocyte (G) UMAP projection split by genotype.

We were primarily interested in fibroblast gene expression profiles in *Yap*^fl/fl^;*Wwtr1*^fl/+^;*Postn^MCM^* hearts following MI compared to control, as this comparison could provide insights into the mechanisms downstream of Yap and Wwtr1 that mediate adverse cardiac remodeling or fibrotic phenotypes. Notably, Yap and Wwtr1 expression was almost exclusively detected in fibroblasts (cluster 5) (**Figure 4D**). Differential expression analysis on fibroblasts (cluster 5) revealed 319 differentially expressed genes (DEGs) between *Yap*^fl/fl^;*Wwtr1*^fl/+^;*Postn^MCM^* and *Postn^MCM^* genotypes (**Table S5**). Upstream regulator analysis of DEGs indicated fibroblasts from *Yap*^fl/fl^;*Wwtr1*^fl/+^;*Postn^MCM^* mice upregulated pathways related to proinflammatory cytokines (IFNγ, STAT1, TNF) and downregulated pathways related to cell cycle activation (Myc (19) and Trim24 (20)) and pro-fibrotic cytokine activation such as IL4 (21) (**Figure 4E**). Amongst the most strongly upregulated genes in fibroblasts from *Yap*^fl/fl^;*Wwtr1*^fl/+^;*Postn^MCM^* hearts included genes associated with collagen secretion and pro-tumorigenic fibroblast properties (*Saa3*, *Mpp6*, *Pdgfra* (22, 23)). Amongst the most downregulated genes included proto-onco genes (*Laptm4b*(24), *Clec3b* (25)) and the myogenic marker, Desmin (*Des* (26)). Of particular interest, the secreted matricellular protein *Ccn3* was amongst the most strongly suppressed in *Yap*^fl/fl^;*Wwtr1*^fl/+^;*Postn^MCM^* fibroblasts (average Log_2_ fold change= −1.69, adjusted p-value 1.24×10^-14^). While CCN family members, *Ccn1* (*Cyr61* (27)) and *Ccn2*, (*Ctgf* (28, 29)) are well known transcriptional targets of Yap and Wwtr1, *Ccn3* expression has not been previously linked to the Hippo-Yap pathway.

To improve resolution of fibroblast phenotypes, we re-clustered cells from fibroblast cluster 5 which resulted in 6 distinct clusters 0-5 (**Figure 4F**). Cells from both *Yap*^fl/fl^;*Wwtr1*^fl/+^;*Postn^MCM^*and *Postn^MCM^* genotypes populated clusters 3 and 5 while cluster 0, 2, and 4 were derived primarily from *Postn^MCM^* hearts and cluster 1 derived primarily from *Yap*^fl/fl^;*Wwtr1*^fl/+^;*Postn^MCM^* hearts (**Figure 4G**). Cluster 0 was highly enriched for anti-proliferative and pro-fibrotic IGF binding proteins (*Igfbp5, Igfbp6 Igfbp3* (30, 31)) as well the epicardial marker *Dmkn* (30, 32–34). Cluster 2 was enriched for genes encoding chemotactic genes (*Cxcl13*, *Cxcl1*, *Cxcl12*, *Apln*), and Cluster 4 was strongly enriched for matrifibrocyte markers (*Angptl7*, *Thbs1*, *Sfrp2*) described by Forte et al as well as *Vegfc* (35). Thus, fibroblasts derived primarily from *Postn^MCM^* control hearts appear to be enriched for pro-fibrotic and pro-inflammatory genes and more closely resemble matrifibrocyte phenotype designed to support a rigid scar (2). Cluster 1 was enriched for epicardial markers (*Saa3*, Mpp6 (35)) suggesting distinct activation of epicardial derived fibroblasts in *Yap*^fl/fl^;*Wwtr1*^fl/+^;*Postn^MCM^* hearts. Cluster 3 contained cells from both genotypes and was strongly enriched for genes described by Forte et al as proliferating myofibroblasts (*Cenpa*, *Hmgb2*, *Cdc20*, *Birc5*, *Cks2*, *Stmn1*, *Top2a*, among others (35)). Cluster 5 contained genes from both genotypes and was strongly enriched for genes encoding ECM components (*Mfap4*, *Col8a1*, *Col14a1*, *Col15a1*) and contractile proteins (*Postn*, *Acta2*, *Tagln*) suggesting a secretory myofibroblast phenotype (36). Collectively, differential gene expression and marker analysis of fibroblast sub-clusters indicates fibroblasts from *Yap*^fl/fl^;*Wwtr1*^fl/+^;*Postn^MCM^* hearts are less proliferative and secretory, and display distinct inflammatory chemokines compared to *Postn^MCM^* hearts.

### Yap and Wwtr1 co-depletion synergistically modulates *Ccn3* gene expression in cardiac fibroblasts

We next sought to address if gene expression changes observed in vivo in *Yap*^fl/fl^;*Wwtr1*^fl/+^;*Postn^MCM^* cells were associated with depletion of either Yap or Wwtr1 individually or required depletion of both factors. We performed siRNA mediated knockdown of *Yap* and/or *Wwtr1* in cultured primary rat neonatal cardiac fibroblasts which resulted in greater than ∼70% depletion of *Yap* and *Wwtr1* as verified by qRT-PCR (**Figure 5A**). Compared to the negative control and single knockdowns of either Yap or Wwtr1, we identified distinct genetic profiles for *Yap*/*Wwtr1* double knockdown cells as evident by a principal component analysis (PCA) of DEGs (**Figure 5B**). A relatively large set of significantly DEGs (p < .05) with Log_2_ fold change (Log2FC) >2 or <-2 were unique to *Yap*/*Wwtr1* double knockdown cells (252 genes) while *Yap* or *Wwtr1* single knockdowns shared the majority of their DEGs (**Figure 5C**), suggesting high functional redundance of Yap and Wwtr1 in cardiac fibroblasts. A large gene cluster that was activated only when both Yap and Wwtr1 were knocked down contained genes primarily related to the immune response (**Figure 5D**). A gene cluster that was significantly downregulated in the Yap/Wwtr1 knockdown group contained genes primarily regulated by Tgfβ-1 and related to “hepatic fibrosis”, Stat3 pathway, and Th2 pathway— illustrating suppression of pro-fibrotic genes with depletion of Yap and Wwtr1. Gene ontology analysis of DEGs unique to Yap and Wwtr1 co-depletion were related to extracellular space (**Figure 5E**) indicating modulation of matrix or secreted proteins. We specifically measured expression of genes that were most strongly differentially expressed in the fibroblast cluster (cluster 5) from our in vivo scRNAseq experiment to determine if their expression was regulated by depletion of Yap, Wwtr1, or both Yap and Wwtr1. Out of the top five downregulated genes from the in vivo scRNAseq experiment, *Ccn3* was the most highly expressed in negative siRNA treated cultured cardiac fibroblasts (**Figure 5F**). Of these genes, only *Ccn3* was robustly suppressed with Yap+Wwtr1 siRNA mediated depletion (**Figure 5G**). While expression of most genes tested showed a synergistic (*Cfb*, *Mgp*, *Prss23*) or additive (*B2m*, *Irf7*, *Isg15*, *Tnfrsf11b*) response to Yap and Wwtr1 depletion, *Anxa2* appeared primarily regulated by Yap while *Ccn3* and *S100a10* more strongly regulated by Wwtr1 (**Figures S9A,B**). From these data we observed a synergistic role between Yap and Wwtr1 regulating gene expression in cardiac fibroblasts, with *Ccn3* expression of particular interest given its prominence in the datasets and function as a matrix element.

**Figure 5.**
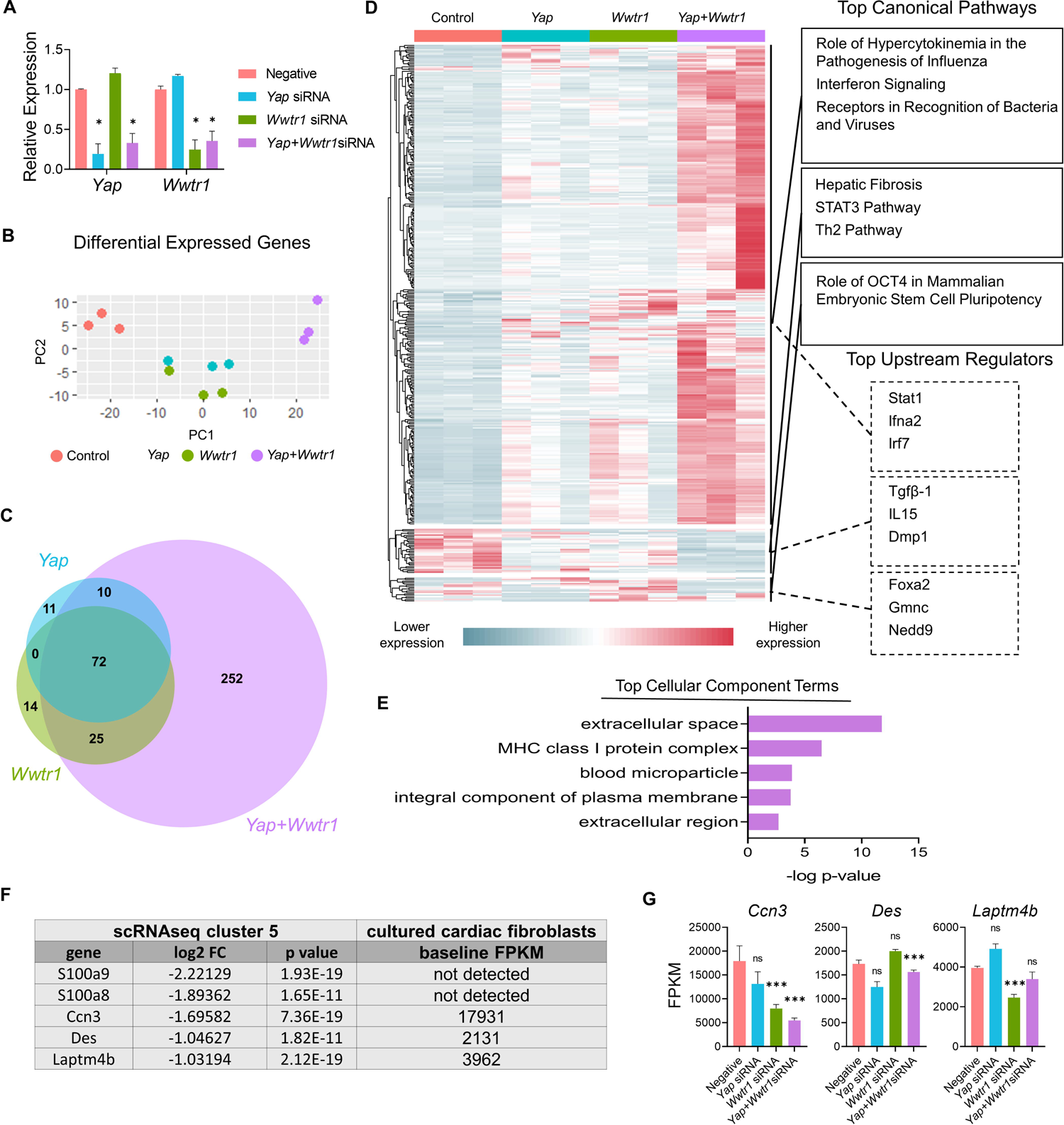
Gene expression following Yap and/or Wwtr1 depletion in vitro. (A) Relative RNA expression of *Yap* and *Wwtr1* after siRNA knockdown in primary cardiac fibroblasts. n = 3 negative control, n = 4 *Yap*, n = 4 *Wwtr1*, n=4 *Yap*+*Wwtr1*. Each data point represents a biological replicate derived from cells from independent pooled litters. One-way ANOVA comparing experimental groups to negative siRNA control by Dunnett’s test. (B) PCA of DEGs from RNAseq from siRNA treated cells n = 3 per group. (C) Venn Diagram denoting unique and common DEGs with pAdj <0.05 and log2FC >2 or <-2 from *Yap*, *Wwtr1, and Yap/Wwtr1* knockdown (D) Heat map showing relative DEGs with pAdj <0.05 and log2FC >2 or <-2. The top three canonical pathways and top three upstream regulators identified for each gene cluster by IPA are listed. (E) The top cellular component terms derived from genes uniquely differentially expressed after Yap/Wwtr1 knockdown. (F) Table depicting the top downregulated differentially expressed genes between *Yap^fl/fl^;Wwtr1^fl/+^;Postn^MCM^* and *Postn^MCM^* fibroblasts and the baseline expression in FPKM values for negative siRNA treated primary cardiac fibroblasts. (G) FPKM values of the genes depicted in (F) following siRNA treatment. ns = not significant, * = P <0.05, and *** = P <0.001.

### CCN3 administration contributes to adverse ventricular remodeling and fibrosis post MI in mice

*Ccn3* was the most strongly downregulated gene in *Yap*^fl/fl^;*Wwtr1*^fl/+^;*Postn^MCM^* fibroblasts at 7 dpi (**Table S5, Figures S10A,B**) that was also significantly and robustly decreased following in vitro knockdown of Yap+Wwtr1 in cardiac fibroblasts, but not Yap knockdown alone (**Figure 5G)**. To confirm our results from scRNAseq and bulk RNAseq of cultured fibroblasts, we repeated the siRNA knockdown experiment in neonatal rat cardiac fibroblasts and measured Ccn3 protein levels. In agreements with our RNAsequencing, knockdown of Yap alone showed no change in Ccn3 protein levels, whereas Yap+Wwtr1 knockdown significantly decreased Ccn3 protein expression (**Figures 6A,B and S11**). Ccn3 is a member of the CCN (*Cyr61, Ctgf, Nov*) family of secreted extracellular proteins (37). While studies have linked Ccn3 to integrin and Notch1 mediated signaling (38) and prevention of renal fibrosis (39), the role of Ccn3 in the heart post injury is virtually unexplored. At the protein level Ccn3 is substantially more abundant in infarcted hearts at 14 dpi compared to uninjured hearts and its expression is decreased in *Yap*^fl/fl^;*Wwtr1*^fl/+^;*Postn^MCM^* compared to *Yap*^fl/fl^;*Wwtr1*^fl/+^ (**Figures 6C and S12**). These data mirror observations in human patients suffering from dilated cardiomyopathies, who show also show elevated cardiac CCN3 expression (40). Furthermore, *Ccn3* expression is strongly enriched in cardiac fibroblasts when compared to other interstitial cells (**Table S6**, **Figures 6D and S13**) suggesting fibroblasts are the primary source of CCN3 in the heart. We hypothesized that *Ccn3* downregulation in *Yap*^fl/fl^;*Wwtr1*^fl/+^;*Postn^MCM^* fibroblasts contributes to improved cardiac functional outcomes post MI, and therefore CCN3 overactivation would promote adverse cardiac remodeling. To test this hypothesis, we performed MI on adult C57/B6 mice and subsequently administered mice with either recombinant CCN3 or vehicle (PBS) by intraperitoneal injection 3 times per week, beginning at 1 dpi. A prior study performed same protocol to investigate the role of CCN3 on fibrosis linked to diabetic nephropathy (39). Strikingly, we found at just 3 dpi that compared to vehicle, mice receiving CCN3 already started to show a decline in cardiac function, and by 14 and 28 dpi cardiac function was substantially and significantly worse (**Figures 6E and S14**). LVID-s increased significantly with administration of CCN3 over the 28-day experiment while LVID-d trended towards being larger (P = 0.08 at 28 dpi, **Figure 6E**). Heart weight and body weight were not significantly different between groups (**Figure 6F**). Histological analysis at 28 dpi revealed significantly larger scars (**Figures 6G,H**) and increased proliferation of scar associated cells in response to CCN3 administration (**Figures 6I,J**) but no difference in cardiomyocyte cross sectional area (**Figure 6K,L**).

**Figure 6.**
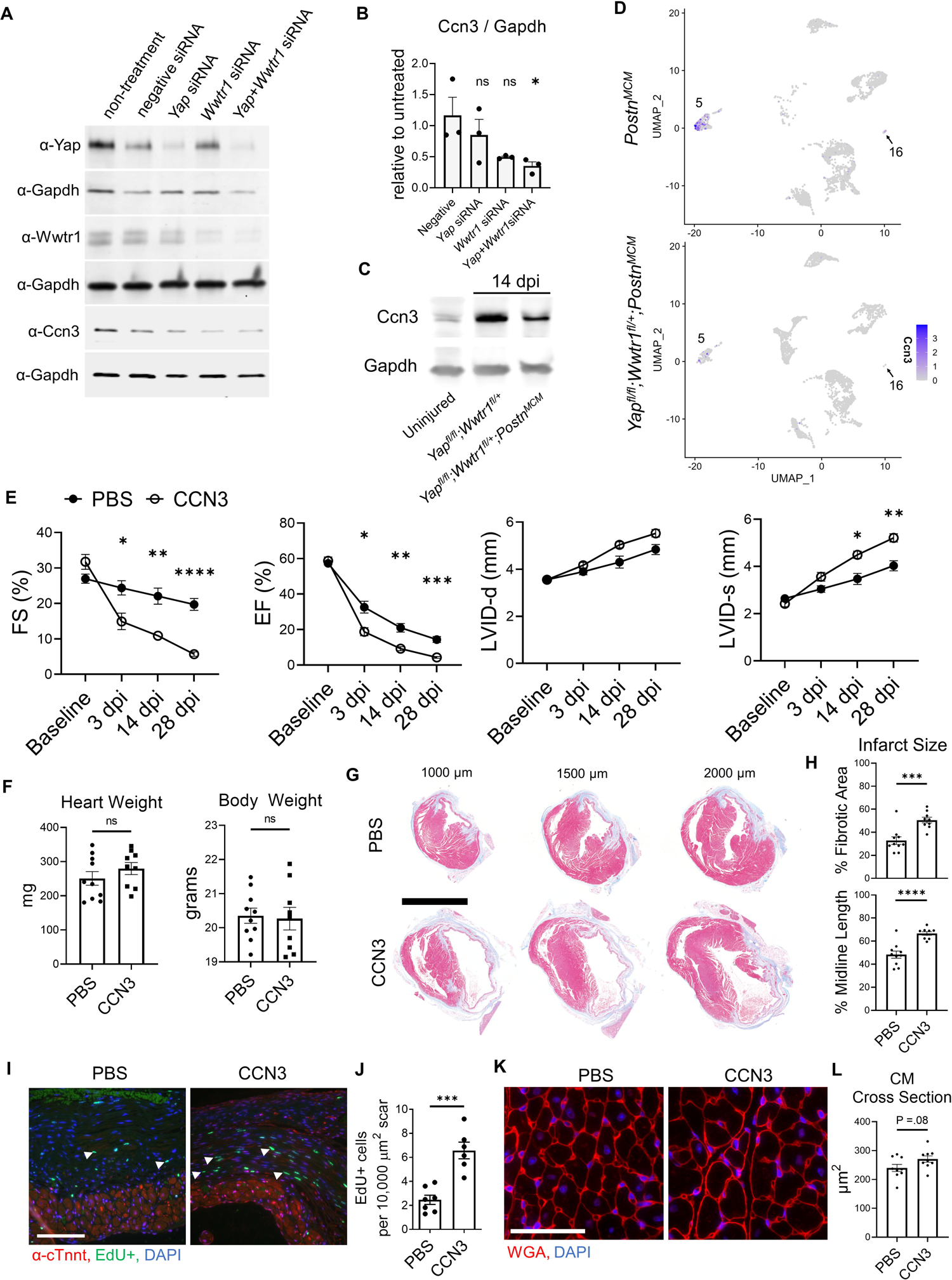
Ccn3 contributes to adverse ventricular remodeling following MI. (A) Western blots depicting protein abundance of Yap, Wwtr1, or Ccn3 following siRNA treatment in cultured neonatal rat cardiac fibroblasts with Gapdh loading controls. (B) Quantification of Ccn3 protein from neonatal rat cardiac fibroblasts following siRNA, normalized to non-treated samples. n = 3 biological replicates per group. Each replicate consisted of cells from independent litters. One-way ANOVA comparing experimental groups to negative siRNA control by Dunnett’s test. (C) Western blot depicting Ccn3 abundance in left ventricles of uninjured and 14 dpi left ventricles. (D) Feature plot illustrating the abundance of *Ccn3* expressing cells by UMI within clusters 5 and 16. (E) Quantification of left ventricular function by fractional shortening, ejection fraction, and left ventricular internal diameters during diastole and systole at baseline. 3, 14, and 28 dpi timepoints were analyzed by repeated measures two-way ANOVA and Sidak multiple comparisons test. n = 10 PBS treated, 9 CCN3 treated. (F) Quantification of heart and body weights at 28 dpi. n = 10 PBS treated, 9 CCN3 treated. Unpaired student’s t-test. (G) Representative serial sections of Gömöri trichrome stained hearts measured as distance from the apex. Scale = 5mm. (H) Quantification of infarct scar size by either total fibrotic area or midline size of the left ventricle. n = 10 PBS treated, 9 CCN3 treated. Unpaired student’s t-test. (I) Representative immunohistological images denoting EdU+ nuclei within the scar area. White arrows indicate EdU+ scar associated nuclei. Scale = 100 µm. (J) Quantification of EdU+ scar associated nuclei. n = 7 PBS treated, 6 CCN3 treated. Scale bar = 100 µm. Unpaired student’s t-test. (K) Representative images and (L) quantification of WGA staining in the remote zone of the left ventricle. n = 9 PBS treated, 9 CCN3 treated. Scale = 100 µm. Unpaired student’s t-test. ns = not significant. F-H are from 28 dpi hearts. ns = not significant, * = P <0.05, ** = P <0.01, *** = P <0.001, **** = P <0.0001.

We next tested assessed the transcriptional changes that occur directly to myocardial tissue following CCN3 administration. We performed MI and administered CCN3 or PBS for 3 consecutive days starting at 1 dpi. Hearts were collected 4 hours after the final injection and left ventricular tissue was processed for RNAsequencing. RNAseq data was normalized to remove unwanted variation, resulting in distinct genetic profiles between CCN3 and PBS control treated animals (**Figure 7A**). We observed extracellular matrix associated genes were predominantly upregulated in ventricles of CCN3 treated mice while transcripts related to mitochondrial function were suppressed (**Figures 7B,C,D**). Thus, we illustrate CCN3 administration following injury drives fibrotic gene networks in myocardial tissue. Further assessment of DEGs by IPA was performed to identify which pathways were modulated. Tgfβ1, a well characterized promoter of fibrosis, was the most activated upstream regulator while Tead1, a transcription factor activated with Yap/Wwtr1 activity, was the most strongly inhibited (**Figure 7F**) (8, 41). Interestingly, while genes mediated by Tead1 activity were predominantly suppressed, indicating a potential repression of Yap/Wwtr1 activity, the expression of the matrix associated genes and CCN family members, *Ccn1* and *Ccn2,* were significantly increased (**Figure 7G**). Ccn1 is promoted by and subsequently drives Tgfβ1 activity, promoting fibrosis in the heart (42, 43) and our data suggests a novel role for exogenously administered CCN3 in contributing to Ccn1, Ccn2, and Tgfβ1 signaling in the heart and aggravating fibrotic remodeling post MI.

**Figure 7.**
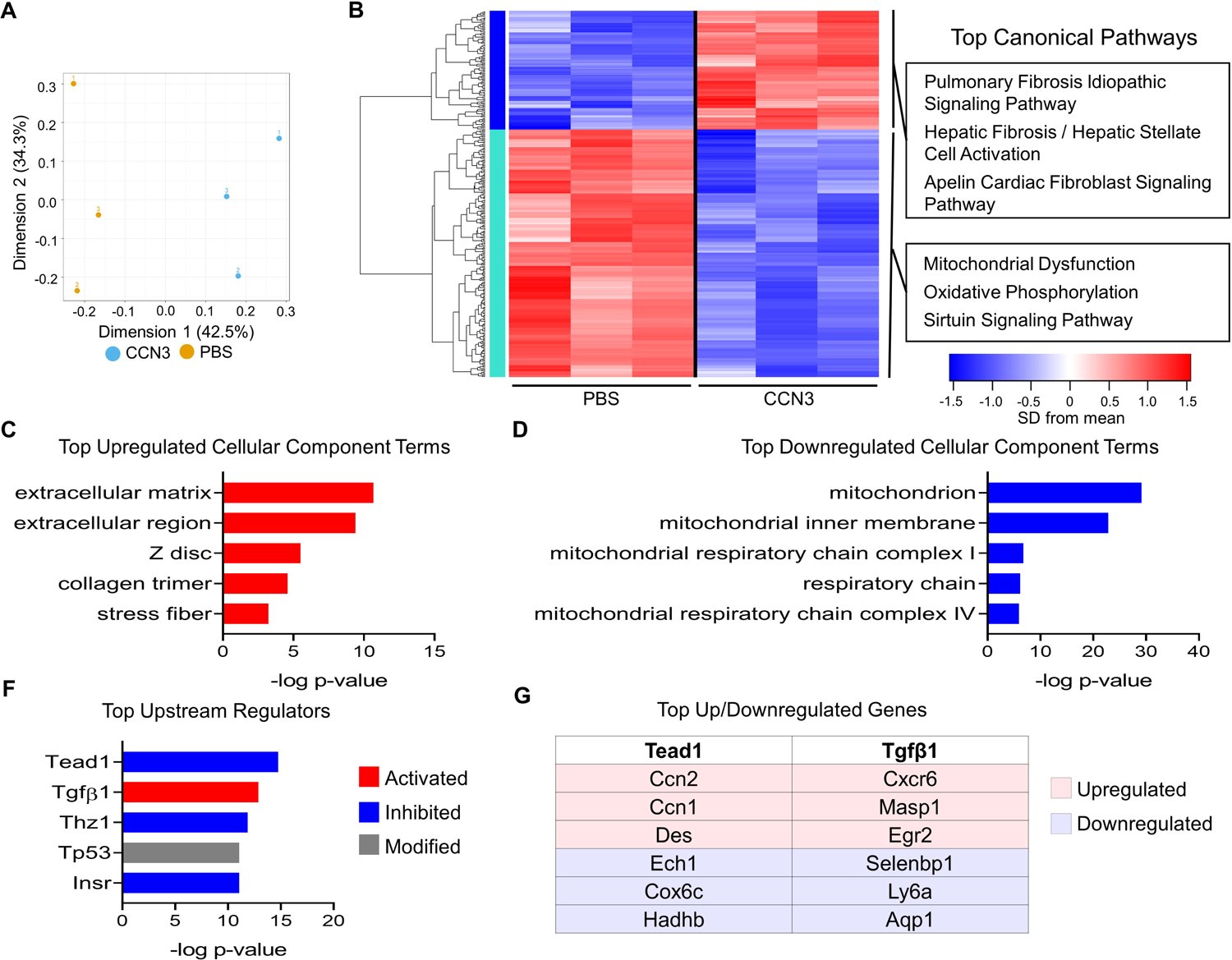
Ccn3 administration promotes fibrotic gene expression post MI. (A) PCA of DEGs from RNAseq from left ventricles. n = 3 per group. (B) Heatmap of 309 genes with one-way ANOVA FDR p-value < 0.2. The top three canonical pathways identified for each gene cluster by IPA are listed. (C and D) The top cellular component terms derived from upregulated (red) or downregulated (blue) DEGs derived from DAVID analysis. (F) The top upstream regulators from all DEGs predicted by IPA. (G) The top upregulated and downregulated genes by fold-change mediated by Tead1 or Tgfβ1.

## Discussion

A nuanced understanding of how myofibroblasts function during wound healing as they proliferate (2), migrate (44), secrete matrix and cytokines (45, 46), recruit immune cells (47), and facilitate a multitude of other roles is salient to understanding the complex nature of progressive heart failure. A therapeutic means to promote the beneficial nature of myofibroblasts (early matrix deposition after injury and recruitment of anti-inflammatory immune cells) while reducing deleterious aspects (latent fibrosis) would be beneficial to curbing heart failure. To this extent, our studies characterize a means by which myofibroblast proliferation and production of cell matrix genes is regulated by the Hippo-Yap pathway.

Although the role of the Hippo-Yap pathway has been studied in progenitor epicardial cells and resident cardiac fibroblasts (11–14), we take the novel approach of assessing the role of endogenous Yap and Wwtr1 expression specifically in myofibroblasts, the cell type responsible for the lion’s share of matrix deposition following MI (1). While deletion of Yap alone did not result in observable changes in scar size or fibrosis, co-disruption of Yap/Wwtr1 resulted in significantly improved cardiac function as well as reduced scar size, interstitial fibrosis, and increased denatured collagen. These results are similar to depletion of Yap and Wwtr1 in resident cardiac fibroblasts whereby Yap and Wwtr1 have been shown to regulate the transition of cardiac fibroblasts to a myofibroblast state (13). However, as therapeutic strategies would likely be implemented after an injury event, once myofibroblast are already activated, our study indicates that inhibiting Yap/Wwtr1 or downstream functional mediators after the transition has already occurred would a reasonable approach.

We highlight the synergistic role of Yap and Wwtr1 in regulating gene expression in cardiac fibroblasts. Our in vivo data demonstrates a significant improvement in cardiac function in *Yap*^fl/fl^;*Wwtr1*^fl/+^;*Postn^MCM^*whereas minimal improvement was observed in *Yap*^fl/fl^*;Postn^MCM^* animals. However, these studies did not include depletion of Wwtr1 alone. Thus, *Yap*^fl/fl^;*Wwtr1*^fl/+^;*Postn^MCM^* phenotypes could be attributed to either Wwtr1 depletion alone or to Yap and Wwtr1 depletion, necessitating further studies to verify this. Our in vitro transcriptomic data elucidated strong synergy in gene regulation with depletion of both Yap and Wwtr1, and enabled us to identify genes differentially expressed in vivo in *Yap*^fl/fl^;*Wwtr1*^fl/+^;*Postn^MCM^* that were regulated by Wwtr1 or Yap+Wwtr1, but not by Yap alone. This cross reference approach helped us prioritize candidate factors that might be mediating the effects of Yap/Wwtr1 in cardiac myofibroblasts.

Our in vitro and in vivo RNAsequencing experiments pointed to candidate genes whose function has been unexplored in the context of cardiac remodeling. Of these genes, *Ccn3* was of particular interest. CCN family members consist of secreted extracellular proteins which have been shown to interact with extracellular matrix components such as Fbln1 and receptors such as integrins and Notch (38). Other members of the CCN family include the well-studied and direct targets of Yap/Wwtr1 mediated transcription Ccn1 and Ccn2 (28, 29). Literature on the interaction between Ccn3 and Hippo-Yap signaling is sparse, but data indicate Ccn3 expression correlates with Yap/Wwtr1 activity. Administration of the Yap/Wwtr1/Tead inhibitor verteporfin decreases *CCN3* expression in cultured human dermal fibroblasts (48) while a decrease in the ratio of transcriptionally suppressed phosphorylated Yap to unphosphorylated Yap correlated with an increase in *Ccn3* during fragmentation of murine ovaries (49). Similar to our findings in injured mice, endomyocardial biopsies from patients with dilated cardiomyopathy show significantly increased expression of CCN3 (40). In mice, Ccn3 knockout mutants display endocardial defects and delayed ventricular septum fusion during development, but are viable as adults (50). Adult knockout mice exhibit cardiomyopathy denoted by hypertrophy and calcification of the septal wall, but not overt ventricular fibrosis (50). To our knowledge, MI studies have not been performed in Ccn3 knockout mice. Overall, the consequence of Ccn3 expression on fibrosis across tissues is not well defined in the literature. While it has been documented in some models that Ccn3 reduces fibrosis via antagonism with Ccn1 and by extension Tgfβ1 signaling (48, 51), this is not always the case (40, 52, 53). Indeed, we illustrate both Tgfβ1 signaling and *Ccn1* expression are both increased in vivo with administration of CCN3. Furthermore, proliferation of interstitial scar cells following cardiac injury was increased. These results mirror those from Lin et al. where CCN3 has been shown to promote DNA synthesis in cultured human skin fibroblasts in the presence of FGF2 (54). However, as systemic administration of CCN3 does affect the renal system and potentially other organ systems and various cell types, we cannot rule out off target effects such as hypertension that may indirectly impact reduced cardiac function we observed in our experiments. Future studies will be aimed at elucidating the how Yap/Wwtr1 modulates Ccn3 expression in cardiac fibroblasts and the collective mechanisms by which Ccn3 contributes adverse cardiac remodeling.

Together, our work illustrates the intrinsic function of Yap and Wwtr1 in myofibroblast activity which promotes fibrosis and deleterious remodeling of the left ventricle after injury. We demonstrate that Ccn3 expression is regulated by Yap and Wwtr1, and CCN3 administration substantially contributes to adverse cardiac function post MI. As such, the Hippo-Yap pathway, Ccn3, or other downstream elements expressed in cardiac myofibroblasts could be attractive targets for modulating adverse remodeling following MI.

## Methods

### Animals

The following mouse lines from The Jackson Laboratory were utilized: stock 027929 *Yap*^fl^, 029645 *Postn^MCM^*, 030532 *Yap*^fl^; *Wwtr1*^fl^, and 004077 *Rosa26-eGFP*^fl^. Sprague Dawley neonatal rats (Charles River Laboratories) were used for cardiac fibroblast cell isolation. Adult mice were euthanized by administration of isoflurane and subsequent thoracotomy, neonatal mice and rats were euthanized by decapitation.

### Isoflurane Administration

During echocardiography, surgical MI, or euthanasia, adult mice were anesthetized via inhalation of 1 to 3% isoflurane vaporized with compressed O2 at a flow rate of 1 L/min.

### Myocardial infarction

Six- to ten-week-old mice were used for adult MI studies. The left anterior descending (LAD) coronary artery was ligated using the “Rapid Method” originally described by Gao et al (55). Briefly, a 1-2 cm incision was made in the skin on the left lateral side of the thorax. The heart was retracted from the thoracic cavity via the 5th intercostal space and a surgical needle with 6-0 prolene suture was inserted under the LAD artery. The vessel was then tied off to create a permanent infarction. The heart was returned to the thorax and the chest wall compressed along the sternal midline to force out excess air. The outer incision of the skin was closed using monofilament nonabsorbable nylon suture. The mouse was removed from anesthesia and returned to a cage to recover under a warming lamp. Immediately following surgery animals were administered 1.5 mg/kg slow-release Buprenorphine and the following day received oral administration of 5 mg/kg Meloxicam.

P6 neonatal MI surgeries were performed as described in Mahmoud et al (56). Briefly, methods were similar to that of adult MI except P6 mice were anesthetized on ice. Following surgery, mice recovered under a heat lamp. When animals were sternal and active, they were returned to their mother.

### Tamoxifen Treatment

Mice of all genotypes were treated with tamoxifen to induce Cre expression and excision of floxed genes following MI. Cre negative controls received identical tamoxifen dosing. For neonatal mice, tamoxifen (Sigma) was dissolved into a mixture of 25% ethanol and 75% sunflower oil at a concentration of 1.5 mg/mL and injected subcutaneously at a dosage of 30 mg/kg. Postnatal day 6 (P6) neonatal mice were administered with tamoxifen 1, 3, and 5 days post injury (dpi). For adult mice, a diet of 0.4 g/kg tamoxifen citrate chow (Envigo) was provided *ad libitum* starting immediately after MI and continued for the duration of the study.

### CCN3 administration

Following MI in 8 week-old C57/B6J mice, recombinant CCN3 (R&D Systems 1640-NV-050) was resuspended in PBS at a concentration of 0.01 µg/µL, and 6 µg/kg was administered by intraperitoneal injection following surgical MI (Tuesday) and subsequently three times weekly (Monday, Wednesday, Friday) as described in Riser et al (39) over the course of 28 days. At this concentration, CCN3 was shown to elicit renal phenotypes but not overt adverse effects in mice (39). PBS was administered by intraperitoneal injection to vehicle control animals.

### Echocardiography

Echocardiograms were obtained from adult mice using a Vevo 770 with an RMV 707 transducer or Vevo 3100 with an MX550D transducer. Scans were taken of the parasternal long axis and short axis at the papillary muscle level in triplicate and measured in a genotype or treatment blinded manner. We calculated ejection fraction (%EF) based on measurements from tracing the circumference of the LV chamber in long axis mode during systole and diastole. Measurements based on internal diameters in short axis mode at the mid-papillary level were used to calculate fractional shortening (%FS), left ventricular internal diameter during diastole (LVID-d), and systole (LVID-s).

### Histological Analysis

Hearts were fixed in 10% formalin for 48 hours at room temperature prior to processing and paraffin embedding. Sectioning was performed starting at the apex and progressing towards the base of the heart in 250 µM steps. 4 µM sections were collected at each level. Gömöri trichrome staining was performed to assess scar size. Slides were scanned using a Super Coolscan 9000 (Nikon) or for higher resolution slides were scanned using a NanoZoomer 2.0-HT (Hamamatsu Photonics K.K.). For scar size quantification, four sections starting at 1000 µm from the apex and proceeding every 500 µm toward the papillary muscles were averaged to quantify fibrotic area and midline percentage using MIquant (57). Interstitial fibrosis was quantified from five representative images in the remote region of the left ventricle and quantified using Fiji ImageJ colorimetric analysis. Immunohistology was evaluated using a Nikon A1 confocal microscope or Eclipse 80i fluorescent microscope (Nikon) and Panda sCMOS camera (PCO). Immunostaining was performed with antibodies described in **Table S1**. Click-iT (Thermo Fisher) 5-ethynyl-2’-deoxyuridine (EdU) staining was performed according to the manufacturer’s recommendations. Two to five regions within the scar were averaged per heart to assess EdU incorporation in non-myocyte scar associated cells. Wheat germ agglutinin (WGA) staining was used to identify the cell surface of cardiomyocytes to assess cross sectional area from five representative images within the remote region of the left ventricle. For denatured collagen assessment, serial sections of the left ventricle containing robust scar regions (∼2000 µm from the apex) were treated with collagen hybridizing peptide with Cy3 conjugate (3Helix) according to the manufacturer’s recommendations. Representative images of the injury were quantified for presence of Cy3 versus scar area. Scale bars are included in one panel for each histological subfigure. All panels within a subfigure are represented by the indicated scale bar.

### siRNA knockdown in cultured rat cardiac fibroblasts

Primary cardiac fibroblasts were isolated from 2-day-old rats using a neonatal heart dissociation kit (Miltenyi Biotec) according to the manufacturer’s protocol, followed by percoll gradient separation. 24 hours after plating, cultured cells were transfected for 48 hours with 25 nM siRNAs designed against Yap and/or Wwtr1, or with universal negative control siRNAs (**Table S2**) as detailed in Flinn, et al (58). Untreated cells were cultured in Dulbecco’s modified eagle medium (DMEM, Life Technologies) supplemented with 7.5% fetal bovine serum without siRNA.

### 3H thymidine incorporation in cultured rat cardiac fibroblasts

Forty-eight hours post siRNA transfection cells were treated with DMEM containing ^3^H thymidine for DNA synthesis quantification. Cells were exposed to ^3^H thymidine treated media for 16 hours, fibroblasts were then washed with PBS and lysed with 0.1 M NaOH + 0.1% Sodium Dodecyl Sulfate (SDS). ^3^H thymidine incorporation was quantified on the LS 6500 MultiPurpose Scintillation Counter (Beckman Coulter).

### Single cell RNA Sequencing (scRNAseq)

To identify differentially expressed genes (DEGs) in cardiac myofibroblasts between *Postn^MCM^* and *Yap*^fl/fl^;*Wwtr1*^fl/+^*;Postn^MCM^* in vivo we performed scRNAseq on interstitial cells from hearts following MI. Both strains carried *R26-eGFP* transgene. MI was performed on 8-10 week old mice as described above and animals were put on tamoxifen citrate chow immediately following surgery. At 7 dpi hearts were extracted and retrograde perfused with 50 mL of 1 mg/mL Collagen type II to digest hearts. Following digestion, atria and valves were removed and ventricular tissue was resuspended in PBS and filtered through a 40 µm cell strainer to remove cardiomyocytes and undigested tissue. Single cells were resuspended in 1 ml of ACK (ammonium-chloride-potassium) lysis buffer (Thermofisher Cat# A1049201) for 2 minutes at room temperature and washed with 2 mL of FACS buffer— 1% fetal bovine serum, 0.1% NaN3 Dulbecco’s Phosphate Buffered Saline (DBPS) without Calcium and Magnesium (Lonza). Cells were resuspended in 4 mL of FACS buffer for cell counting, which was performed using the LUNA^TM^ automated cell counter (Logos Biosystems) using 0.4% trypan blue (Thermofisher Cat# T10282) and the LUNA^TM^ cell counter bright field feature. Subsequently, samples were resuspended to a concentration of one million cells per 100 μL of FACS buffer. Samples were stained with 3 μM of (4’,6-Diamidino-2-Phenylindole, Dilactate (DAPI, Biolegend Cat# 422801)), in a volume of 1 mL of DAPI-FACS buffer per sample for 15 minutes. Then samples were washed with 1 mL of FACS buffer and resuspended again in 1 mL of FACS buffer for cell sorting. Samples were sorted using BD FACS Aria II Cell Sorter (BD Bioscences) and collected in 3 to 5 mL of FACS buffer.

Single cell capture, cDNA synthesis, barcoding, and library preparation was performed using the 10x Chromium system using V3.1 chemistry according to the manufacturer’s recommendation (10x Genomics). Each sample was loaded onto a single lane of a Chromium Next GEM chip G to target 3000 cells per sample. Cells were captured in single GEMs and lysed followed by cDNA synthesis using 12 amplification cycles, followed by library construction per manufacturer’s protocol. An i7 multiplex single index kit was used to generate the libraries over 14 cycles of sample index PCR. Fragment size of cDNA and libraries was assessed using Agilent’s 5200 Fragment Analyzer System.

### scRNAseq data analysis

Libraries were sequenced at the Roy J. Carver Biotechnology Center at the University of Illinois, Urbana Champaign on a NovaSeq 6000 using one S4 lane with 2X150nt reads. Samples were demultiplexed using Cell Ranger v6.1.1 (10X Genomics). A custom reference was made using NCBI’s GRCm39 genome and Annotation Release 109, along with Cloning vector pEGFP-1 (GenBank: U55761.1) and SA-betageo synthetic construct (full details of modifications in Supplemental R file, (59)). The ‘cellranger count’ pipeline with default parameters was run separately on each sample to call cells and collapse reads to unique molecular identifiers (UMIs). Both samples were combined using ‘cellranger aggr’ with ‘--normalize=non’.

The UMI counts per gene for all called cells were read into R (v4.1.2, (60)) and analyzed using Seurat (v4.0.6, (61)). Genes were filtered out if they did not have at least 1 UMI in at least 20 cells, leaving 16,855 genes. Initial quality control involved performing sctranform normalization (62), principal components analysis, shared nearest neighbor cluster calling, and Uniform Manifold Approximation and Projection (UMAP, (63)) dimension reduction (hereafter referred to as the “Seurat pipeline”). One cluster of cells had extremely high percentage of UMIs in mitochondrial genes (likely dead/dying cells) and two other clusters had extremely low numbers of genes detected and total number of UMIs (likely stripped nuclei). These 3 clusters were removed completely along with a few other cells that had total numbers of UMIs > 54,742 or percentage of UMIs in mitochondrial genes > 4.58 (thresholds set at 6 median absolute deviations). The remaining cells were re-run through the Seurat pipeline to create the final clustering and dimension reduction.

Marker genes per cluster were found by recursively comparing each cluster’s cells against all other clusters combined using a Wilcoxon Rank Sum test. Within each cluster, gene expression differences between Yap^fl/fl^Wwtr1^fl/+^;Postn^MCM^ and Postn^MCM^ cells were tested also using a Wilcoxon Rank Sum test. The cells that expressed Postn were overwhelmingly in one cluster, so this one cluster was run by itself through the Seurat pipeline to find sub-clusters of cells. Sub-cluster marker genes and within-subcluster Yap^fl/fl^Wwtr1^fl/+^;Postn^MCM^ and Postn^MCM^ DEGs were calculated as before. Full R codes for all Seurat analyses are in the Supple Methods.

### Western blot

Cultured rat cardiac fibroblasts or mouse left ventricles were collected in RIPA buffer. Rat cardiac fibroblasts were collected 48 hours post siRNA transfection. Protein lysates were combined with Laemmli buffer and separated on a 4-15% Mini-PROTEAN TGC precast gel (Bio-Rad) by electrophoresis. Proteins were then transferred to a 0.45µm pore size nitrocellulose membrane (Bio-Rad). Western blots were processed according to Li-Cor’s Near-Infrared Western Blot Detection protocol and blocked using TBS based Intercept buffer (Li-Cor Biosciences). Protein detection was performed using an Odyssey-CLx infrared imager (Li-Cor Biosciences). Uncropped blots are provided in the supplemental data.

### RNA extraction and qRT-PCR

Twenty-four hours post siRNA transfection media was changed to DMEM with 7.5% FBS, and forty-eight hours later cells were collected for gene expression analysis by either qRT-PCR or RNAseq. Cells were washed in PBS and collected in TRIzol for RNA extraction according to the manufacturer’s recommendations. For qRT-PCR analysis RNA was reverse transcribed using a high-capacity cDNA reverse transcription kit (Applied Biosystems). qRT-PCR was performed with SybrGreen (Invitrogen) and primers listed in **Table S3** and amplification was detected on the QuantStudio 6 Flex (Thermo Fisher). Gene expression was normalized to *18s* ribosomal RNA expression and analyzed in QuantStudio (Thermo Fisher).

### Bulk RNAsequencing

Two methods were used to attain bulk RNAseq data. First, primary cardiac fibroblasts were isolated from 2-day-old rats, treated with siRNA or a universal negative control, and RNA was extracted from cells using TRIzol (Thermo Fisher) extraction protocol according to the manufacturer’s recommendations. This process was repeated with separate litters to achieve 3 biological replicates per group. RNA quality was determined using an Agilent BioAnalyzer. RNA libraries were prepared by BGI Americas. Sequencing was performed using a DNBSEQ-G400 platform at 20M reads per sample. Adapter sequences were removed from the output sequence and reads with low base quality (<13) were further trimmed using Trim_Galore v0.6.5 (Babraham Bioinformatics). Trimmed reads were then aligned to the rat genome (rn6) using Hisat2 v 2 2.1. Transcripts were assembled from RNA-seq alignments using Stringtie2 v2.1.5. Expression was quantified by fragments per kilobase of transcript per million reads mapped (FPKM). DEGs for each experimental group, as compared to the negative control, were detected using DESeq2.

Second, left ventricles were obtained from 4 dpi adult mice treated with either 6 µg/kg CCN3 or PBS daily starting at 1 dpi. RNA qas extracted from homogenized tissue by TRIzol extraction. RNA libraries were prepared by the Roy J. Carver Biotechnology Center at the University of Illinois with the Kapa Hyper Standed mRNA library kit (Roche) and sequenced with a NovaSeq 6000 with V1.5 sequencing kits. Fastq files were generated and demultiplexed with the bcl2fastq v2.20 Conversion Software (Illumina). Salmon version 1.4.0 was used to quasi-map reads to the GRCm39 transcriptome (NCBI) and quantify the abundance of each transcript. Data was normalized by removing unwanted variation by a factor of 2 (64). Differential gene expression analysis was performed using the edgeR-quasi method using a model of treatment + 2 RUV factors plus False Discovery Rate (FDR) correction on the p-values.

For both methods, analysis using the Ingenuity Pathway Analysis (IPA, Qiagen) and Database for Annotation, Visualization and Integrated Discovery (DAVID, (65)) was performed on DEGs.

### Data Availability

Data from the bulk in vitro RNAseq, bulk left ventricular RNAseq, and scRNAseq have been uploaded to the Gene Expression Omnibus (https://www.ncbi.nlm.nih.gov/geo/) and can be accessed under GSE185368, GSE217925, and GSE204712 respectively.

### Statistics

Data were analyzed using Prism 8.2.0 (GraphPad). Two-way ANOVA followed by Tukey’s multiple comparisons tests were performed on samples with two experimental factors. For data series consisting of two experimental factors assessed at multiple timepoints, a repeated measures Two-way ANOVA was performed followed by Sidak multiple comparisons test. Statistical comparisons between two groups were analyzed by Student’s t-test, or between three or more groups by one-way ANOVA followed by Tukey’s or Dunnett’s multiple comparisons test. Assessment of survival curves was performed using a Logrank Mantel-Cox test. Error bars in graphical data represent standard error.

### Study Approval

All protocols in these studies were approved by the local Animal Care and Use Committee and conform to the Guide for the Care and Use of Laboratory Animals published by the National Institutes of Health.

## Supporting information

Supplemental_Methods_codes

## Author Contributions

Conceptualization: C.C.O., M.A.F, M.P., B.A.L.; Methodology: M.A.F., S.A-A., J.D., X.Z.; Investigation: M.A.F., S.A-A., J.D., M.C.K., V.A.A., S.J.P., X.Z., T.B., A.J.; Formal analysis: M.A.F., S.A-A., J.D., M.C.K., V.A.A., S.J.P., X.Z., T.B., A.J.; Writing - original draft: M.A.F., C.C.O., S.A-A, J.D.; Writing - review & editing: M.A.F., S.A-A., M.C.K., V.A.A., S.J.P., X.Z., T.B., A.J., P.L., J.D., M.P., B.A.L., C.C.O.; Supervision: C.C.O., M.P., B.A.L., P.L.; Project administration: C.C.O.; Funding acquisition: C.C.O, B.A.L., M.A.F., M.P., S.J.P.

## Acknowledgements

This work was supported by Advancing a Healthier Wisconsin Co-Investigator Grant (B.A.L. and C.C.O.); by the National Institutes of Health (R01 HL141159 to C.C.O., R01 HL155085 to M.P., T32 HL134643 and F32 HL150958 to M.A.F., and F31 HL150919 to S.J.P.); by the Cardiovascular Center’s A.O. Smith Fellowship Scholars Program (M.A.F.); by the American Heart Association (18CDA34110240 to M.P.), and by the Medical College of Wisconsin Cardiovascular Center (FP00012308). We thank the Roy J. Carver Biotechnology Center at the University of Illinois, Urbana Champaign for sequencing and bioinformatic support.

