## Supplemental Data for "Myofibroblast Ccn3 is regulated by Yap and Wwtr1 and contributes to adverse cardiac outcomes"

**Myofibroblast depletion of Yap and Wwtr1 suppresses Ccn3 expression and improves cardiac function after myocardial infarction**

**Supplementary Materials**

Michael A. Flinn PhD, Santiago Alvarez-Argote MD, Makenna C. Knas, Victor Alencar Almeida, Samantha J. Paddock PhD, Xiaoxu Zhou, Tyler Buddell PhD, Ayana Jamal, Pengyuan Liu PhD, Jenny Drnevich PhD, Michaela Patterson PhD, Brian A. Link PhD, Caitlin C. O’Meara PhD

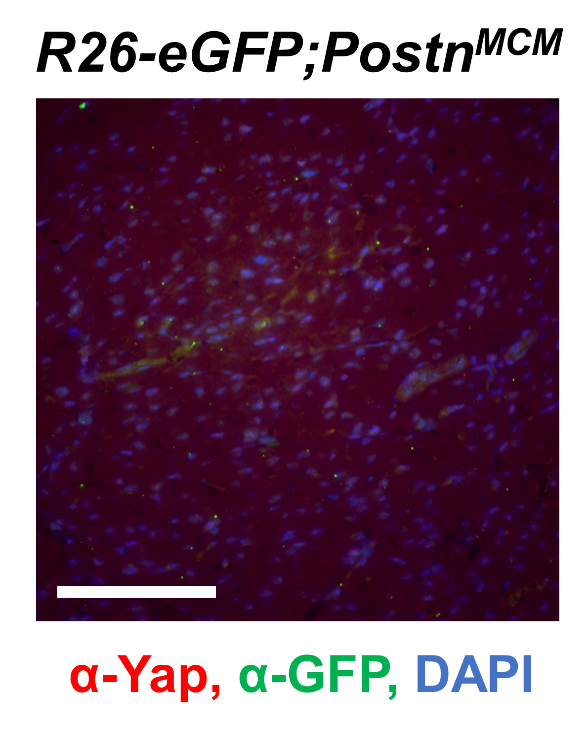

**Figure S1. Periostin expression in remote zone.** Representative image of the remote region in *R26-eGFP^f/+^;Postn^MCM^* littermates. Scale bar = 100 µm.

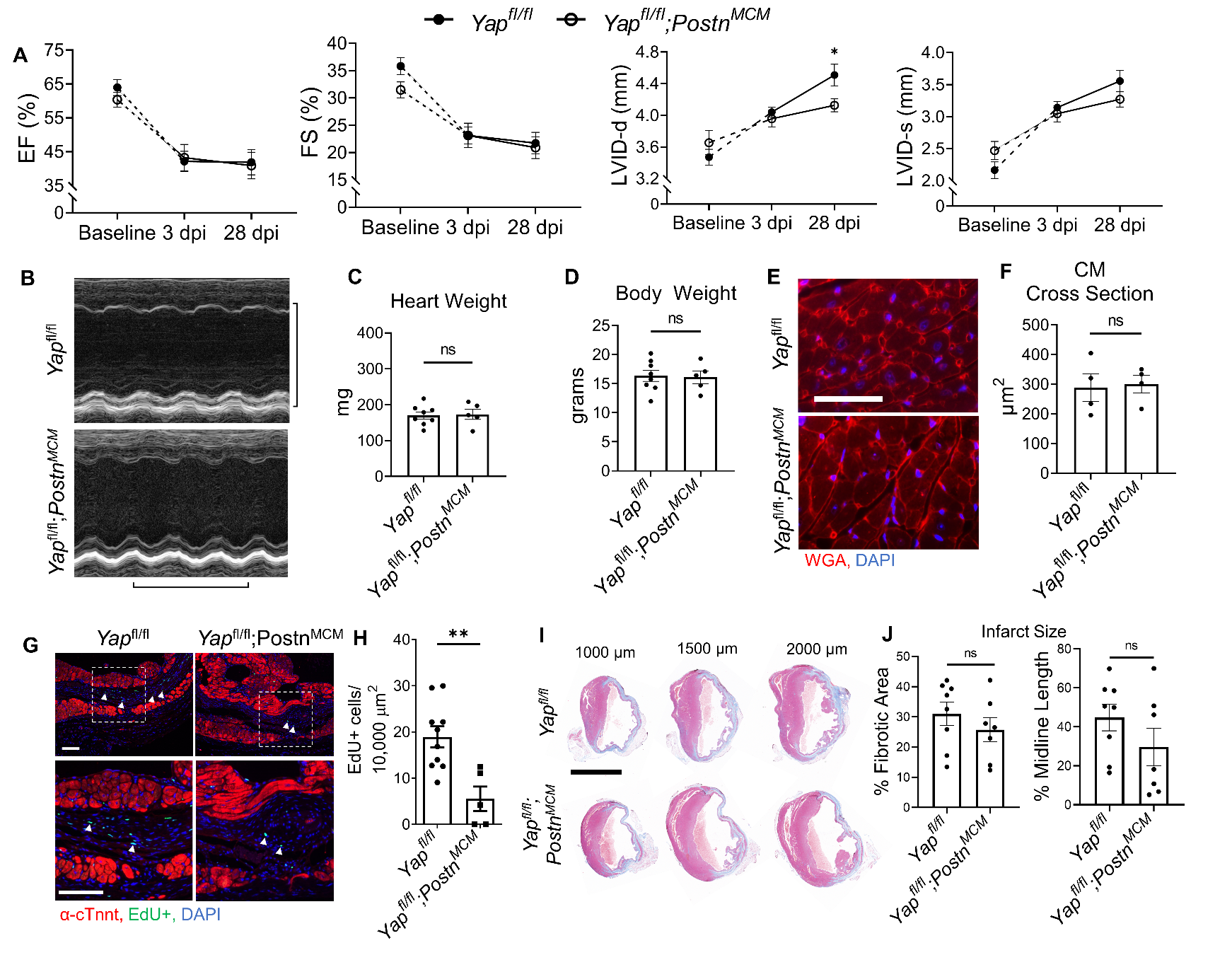
**Figure S2. Deletion of *Yap* from myofibroblasts results in a modest protection against ventricular dilation after injury.** (A) Quantification of %FS, %EF, and left ventricular internal diameters (LVID) during diastole and systole at 3 and 28 dpi analyzed by repeated measures two-way ANOVA and Sidak multiple comparisons test. n = 19 *Yap^fl/fl^* and 13 *Yap^fl/fl^;Postn^MCM^*. A separate subset of animals establishing a baseline as denoted by the hashed lines and assessed by un-paired student’s t-test. n = 11 *Yap^fl/fl^* and 6 *Yap^fl/fl^;Postn^MCM^*. (B) Representative M-mode echocardiograms of left ventricles 28 dpi. Short axis view at mid papillary muscle. Horizontal bar = 500 ms. Vertical bar = 5 mm. (C,D) Quantification of heart and body weights at 28 dpi. n = 8 *Yap^fl/fl^* and 5 *Yap^fl/fl^;Postn^MCM^*. Unpaired student’s t-test. Histological analysis (E-H) was done on 28 dpi hearts. (E) Representative images of WGA staining in the remote zone of the left ventricle. Scale = 50 µm. (F) Quantification of cardiomyocyte cross sectional area. n = 4 *Yap^fl/fl^* and 3 *Yap^fl/fl^;Postn^MCM^*. Unpaired student’s t-test. (G) Representative images of EdU and cardiac troponin (cTnnt) staining within the scar area (scale = 50 µm). White arrows indicate EdU+ scar associated nuclei. (H) Quantification of EdU+ scar associated nuclei. n = 10 *Yap^fl/fl^* and 5 *Yap^fl/fl^;Postn^MCM^*. Unpaired student’s t-test. (I) Representative serial sections of Gömöri trichrome stained hearts measured from the apex. Scale = 5mm. (J) Quantification of infarct scar size by either total fibrotic area or midline size of the left ventricle. n = 8 *Yap^fl/fl^* and 7 *Yap^fl/fl^;Postn^MCM^*. Unpaired student’s t-test. ns = not significant, * = P <0.05, and ** = P <0.01.

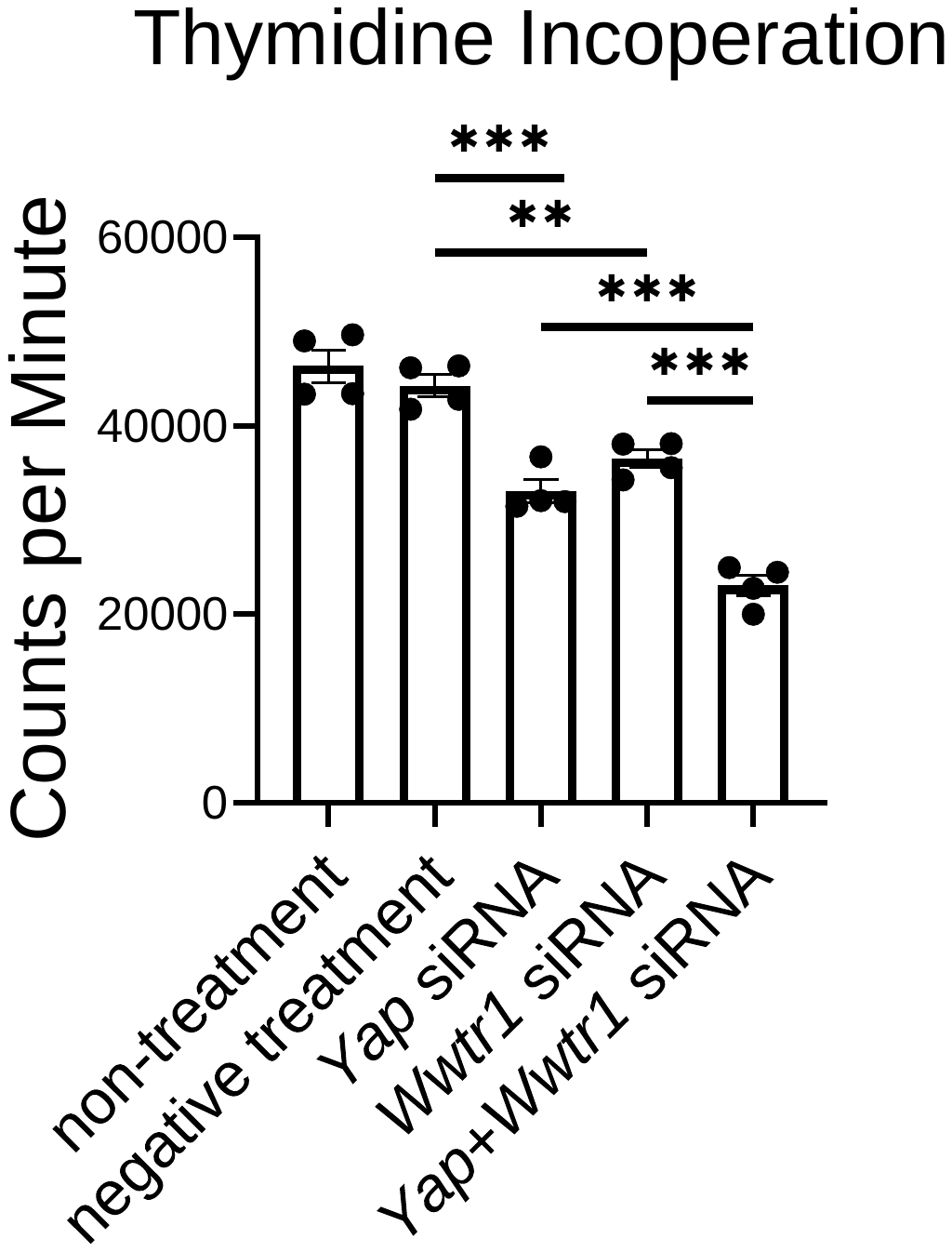

**Figure S3. Yap and Wwtr1 promote cardiac fibroblast DNA synthesis.** Quantification of ^3^H thymidine incorporation in neonatal rat cardiac fibroblasts following siRNA transfection. n = 4. One-way ANOVA, Tukey’s multiple comparisons test. ** = P <0.01, and *** = P <0.001.

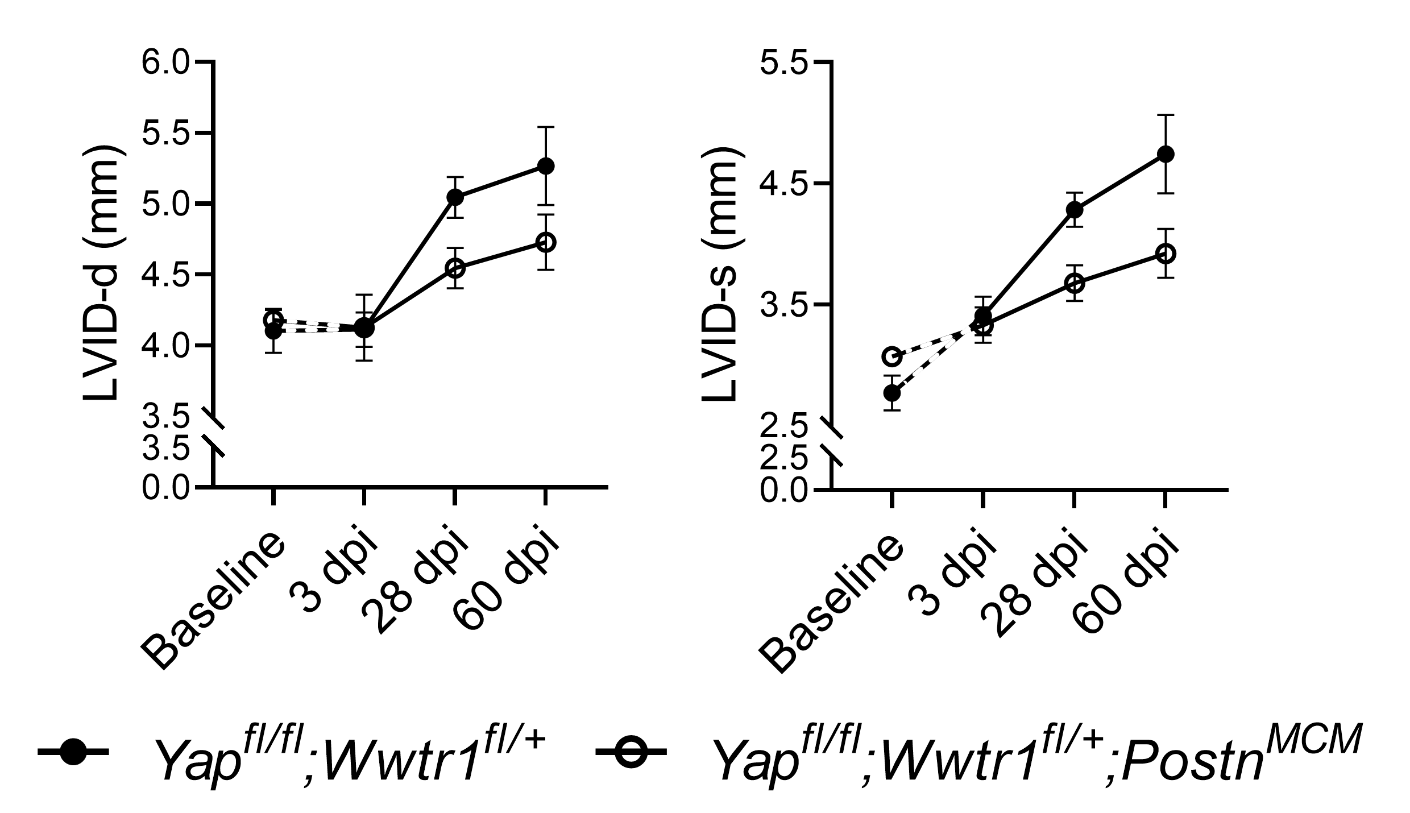

**Figure S4. Left ventricular internal diameters in *Yap^fl/fl^;Wwtr1^fl/+^* and *Yap^fl/fl^;Wwtr1^fl/+^;Postn^MCM^* mice post MI.** Quantification of left ventricular diameter in diastole or systole analyzed by repeated measures two-way ANOVA and Sidak multiple comparisons test. n = 7 *Yap^fl/fl^;Wwtr1^fl/+^*, 4 *Yap^fl/fl^;Wwtr1^fl/+^;Postn^MCM^*. A separate subset of animals establishing a baseline for *Yap^fl/fl^;Wwtr1^fl/+^* animals as denoted by the hashed lines and assessed by un-paired student’s t-test. n = 4 *Yap^fl/fl^;Wwtr1^fl/+^* and 5 *Yap^fl/fl^;Wwtr1^fl/+^;Postn^MCM^*. No data points between genotypes were significantly different at any given timepoint.

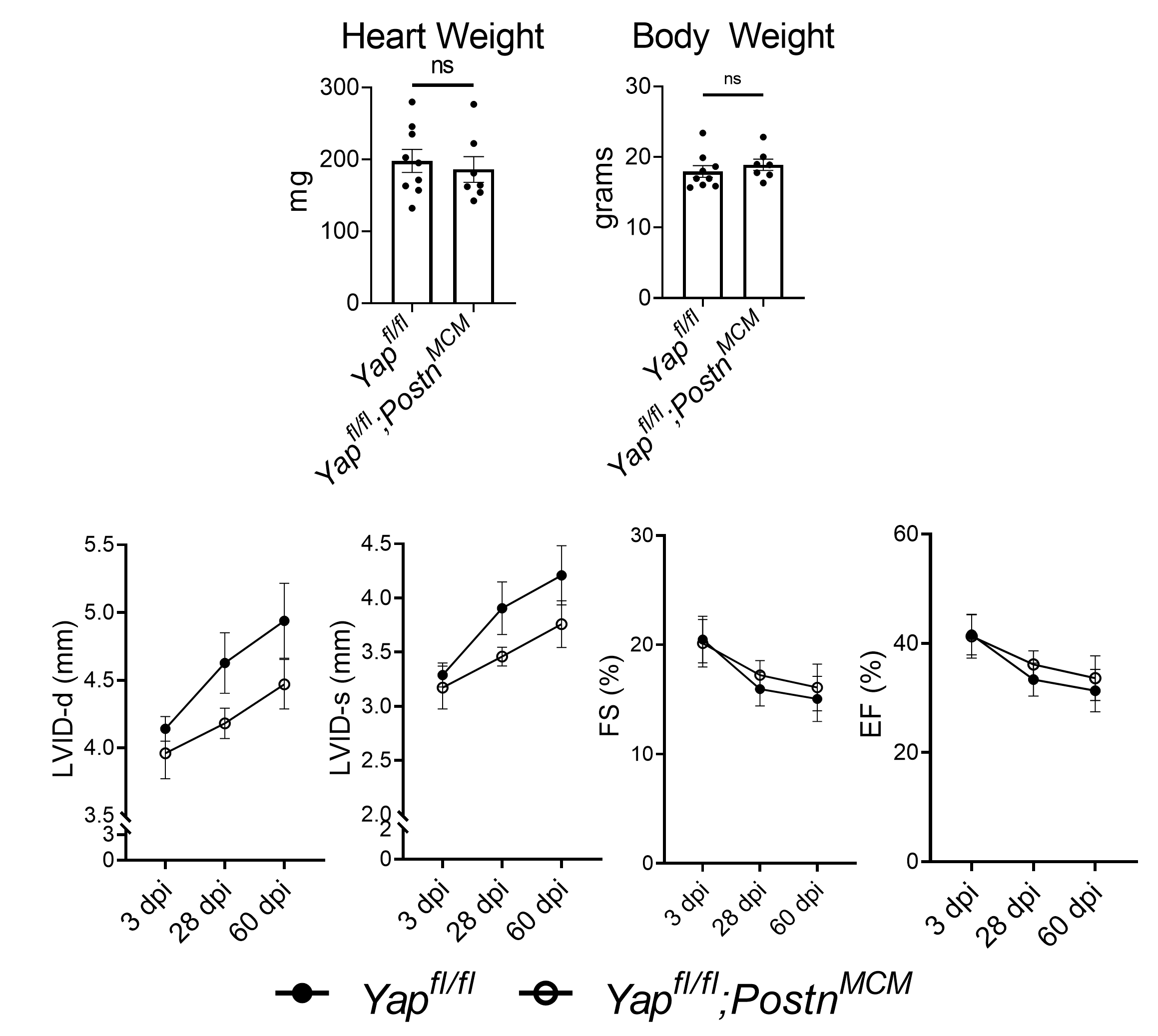

**Figure S5. Cardiac function of *Yap^fl/fl^* and *Yap^fl/fl^;Postn^MCM^* up to 60 days post MI.** Quantification of heart and body weights at 60 dpi. n = 9 *Yap^fl/fl^* and 7 *Yap^fl/fl^;Postn^MCM^*. Unpaired student’s t-test. Quantification of left ventricular function at 3, 28, and 60 dpi analyzed by repeated measures two-way ANOVA and Sidak multiple comparisons test. n = 9 *Yap^fl/fl^;Wwtr1^fl/+^* and 7 *Yap^fl/fl^;Wwtr1^fl/+^;Postn^MCM^*. ns = not significant

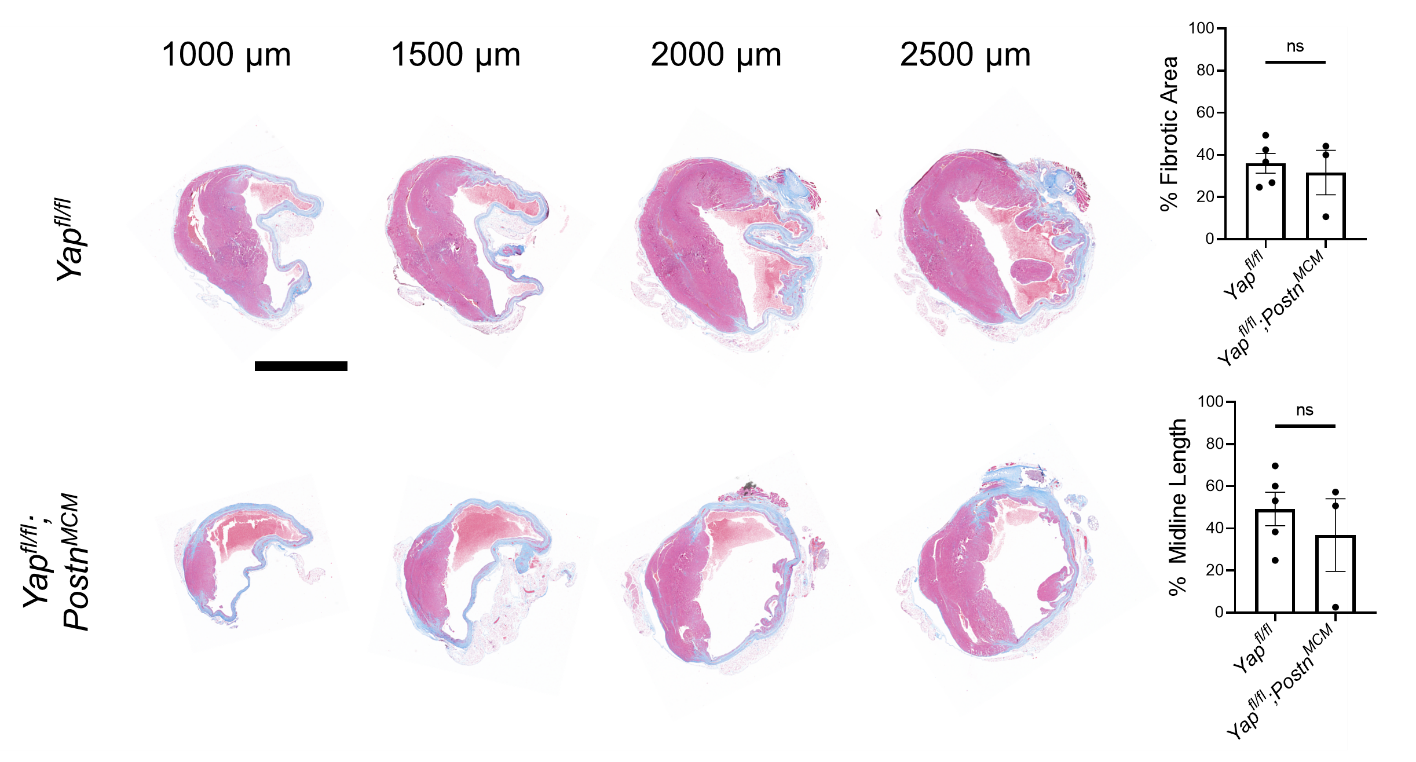

**Figure S6. Scar size at 60 dpi in *Yap^fl/fl^* and *Yap^fl/fl^* *;Postn^MCM^* mice.** Representative serial sections of Gömöri trichrome stained hearts, measured from the apex. Scale = 5mm. Quantification of infarct scar size by either total fibrotic area or midline size of the left ventricle. n = 5 *Yap^fl/fl^* and 3 *Yap^fl/fl^* *;Postn^MCM^*. Unpaired student’s t-test. ns = not significant.

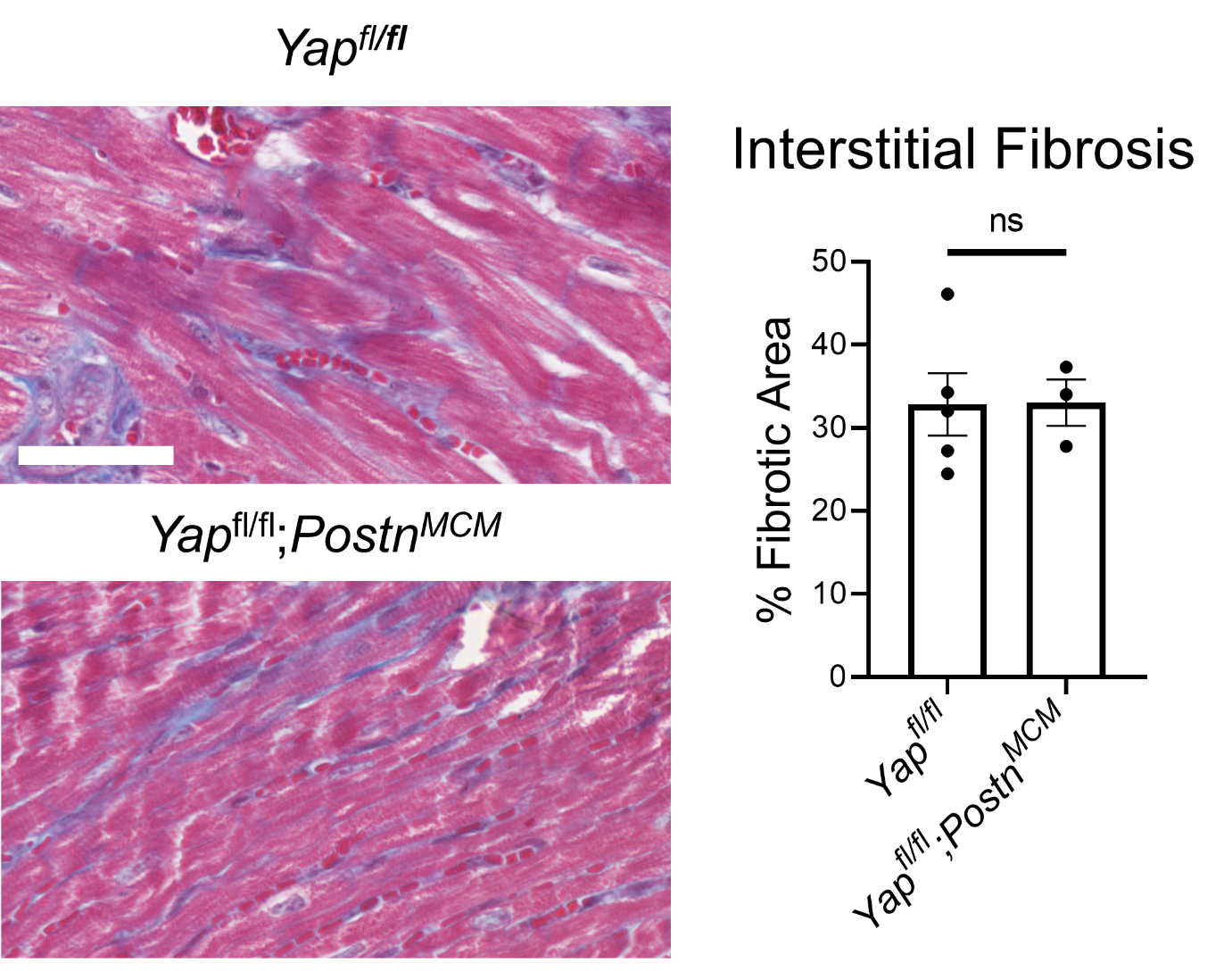

**Figure S7. Myofibroblast knockout of Yap does not alter interstitial fibrosis.** Representative images Gömöri trichrome stained 60 dpi hearts. Sections depict interstitial fibrosis in the remote zone of injured hearts (scale = 50 µm). Quantification of the % blue fibrotic area versus the total left ventricle area. n = 5 *Yap^fl/fl^* and 3 *Yap^fl/fl^* *;Postn^MCM^*. Unpaired student’s t-test. ns = not significant.

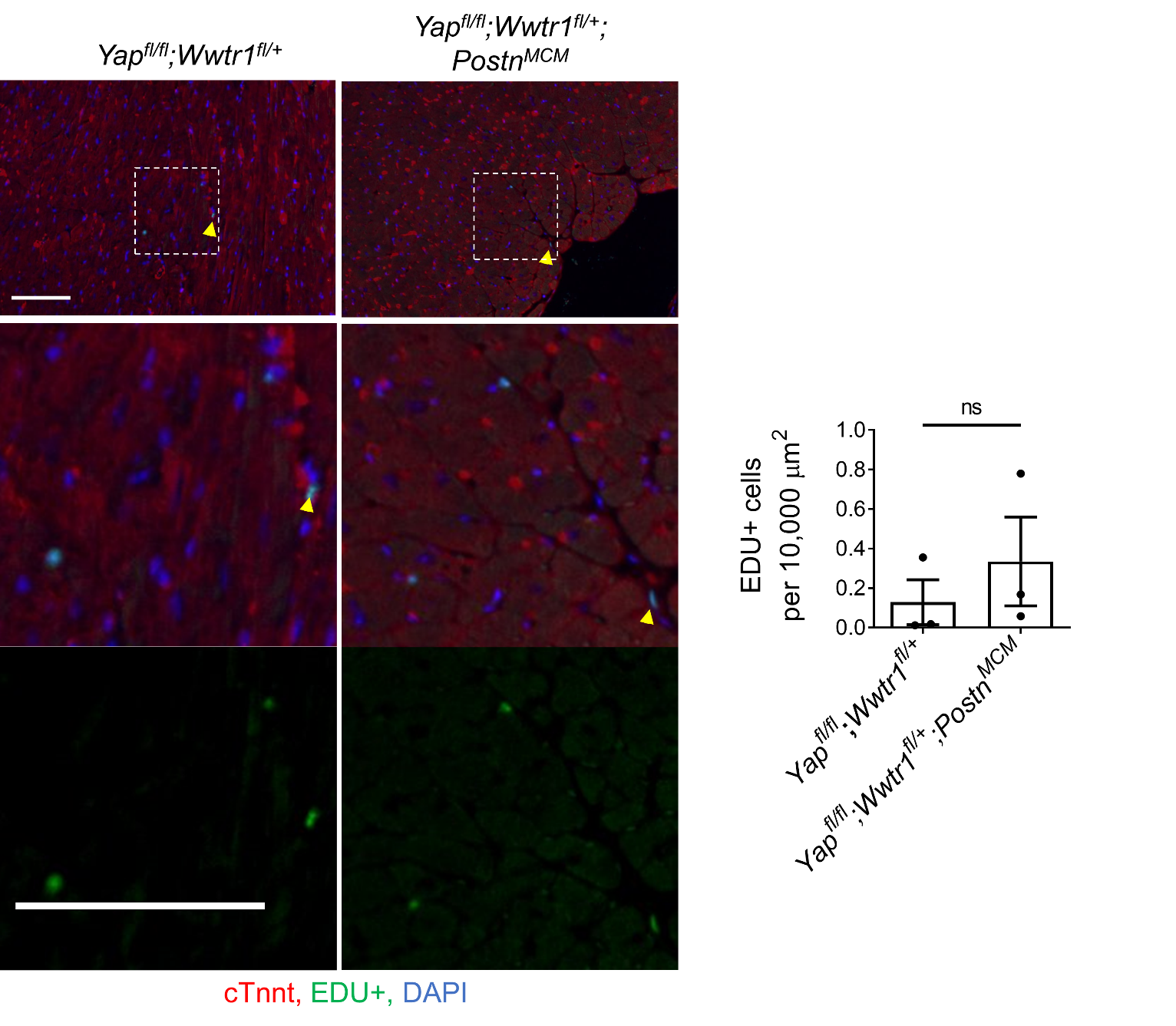

**Figure S8. Remote zone DNA synthesis.** Representative images of EdU incorporation in the remote zone of *Yap^fl/fl^;Wwtr1^fl/+^;Postn^MCM^* and *Yap^fl/fl^;Wwtr1^fl/+^* littermates. Scale = 100 µm. Yellow arrows indicate EdU+ interstitial nuclei. Quantification of EdU+ interstitial nuclei. n = 3 per group. Unpaired student’s t-test. ns = not significant.

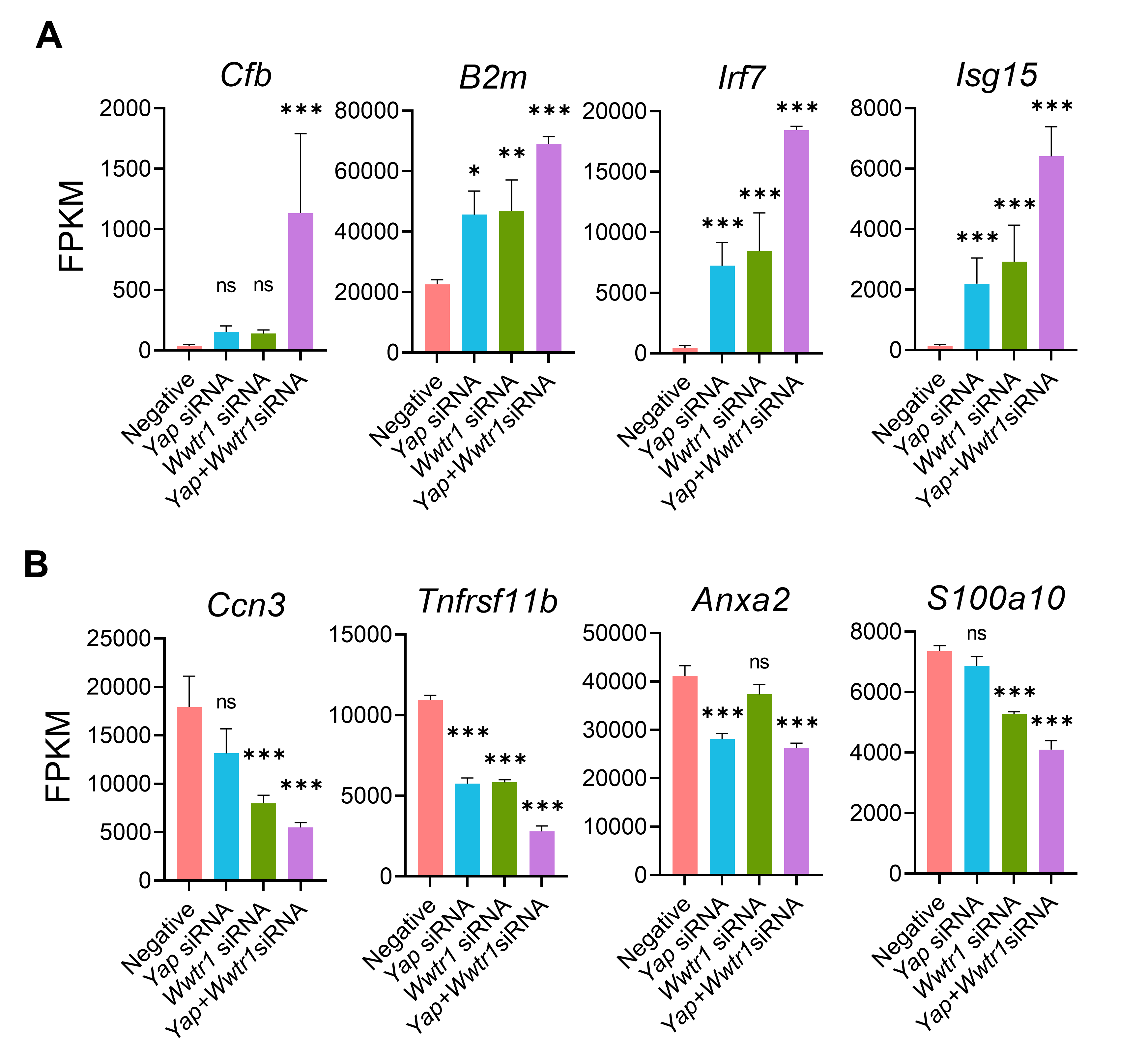

**Figure S9.** FPKM values for top up (A) or down (B) differentially expressed genes between *Yap^fl/fl^;Wwtr1^fl/+^;Postn^MCM^* and *Postn^MCM^* fibroblasts (cluster 5) that also showed the same expression pattern following Yap and/or Wwtr1 siRNA knockdown in vitro. ns = not significant and *** = P <0.001.

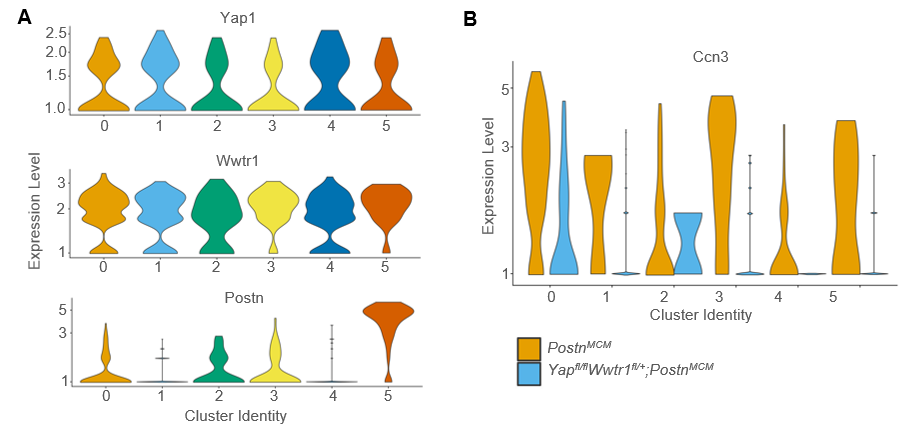

**Figure S10. Cardiac fibroblast subcluster gene expression.** (A) Violin plots denoting UMI count for *Yap*, *Wwtr1*, or *Postn* among each fibroblast subcluster. (B) Violin plots denoting UMI count for *Ccn3* within each subcluster, split by genotype.

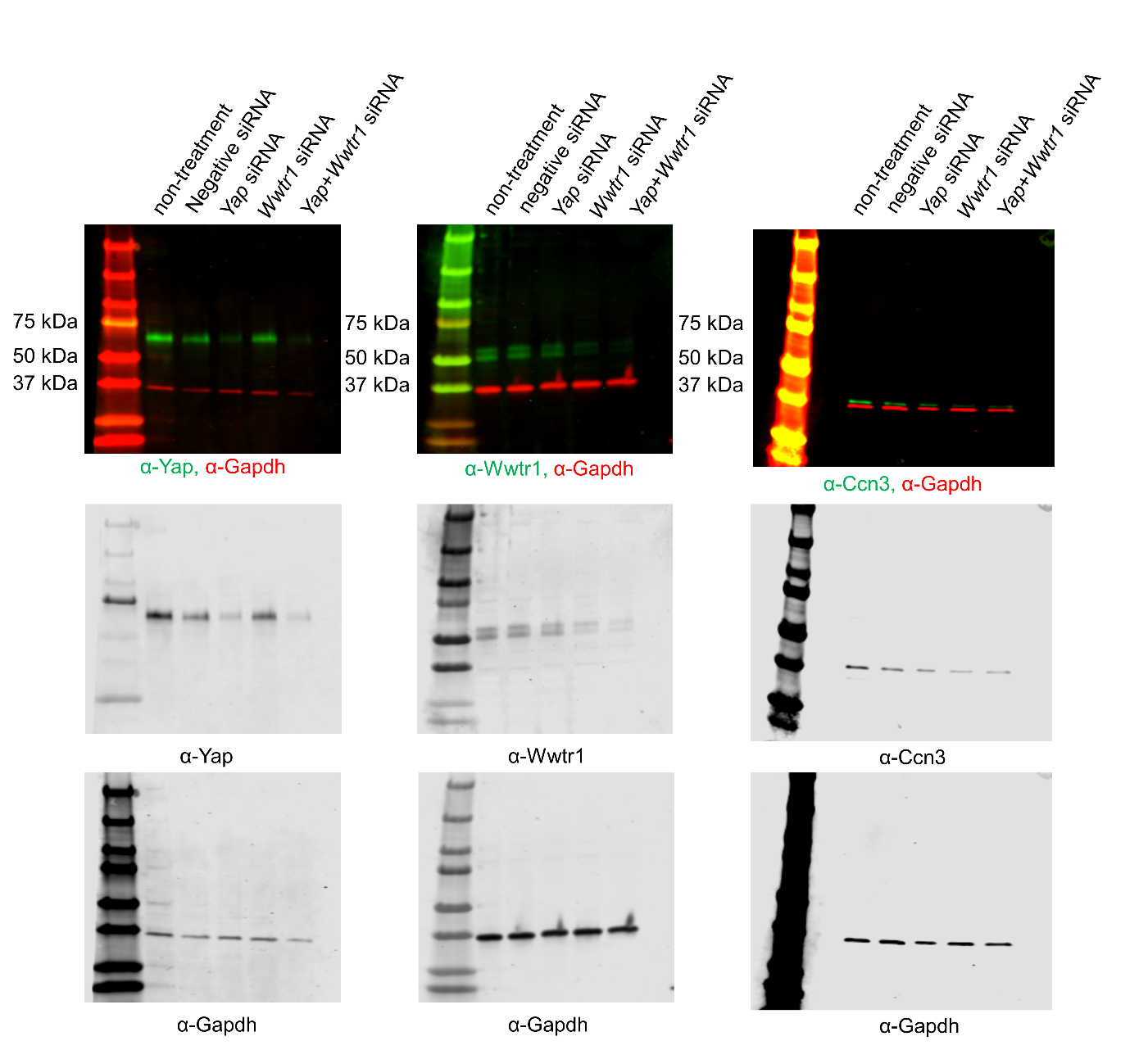

**Figure S11. Ccn3 expression in cardiac fibroblasts with *Yap* and/or *Wwtr1* depletion.** Representative full length western blots illustrating expression of Yap, Wwtr1, Ccn3 and Gapdh in cultured rat cardiac fibroblasts depleted for *Yap* and/or *Wwtr1*. Data presented in Figure 6H are derived from these blots.

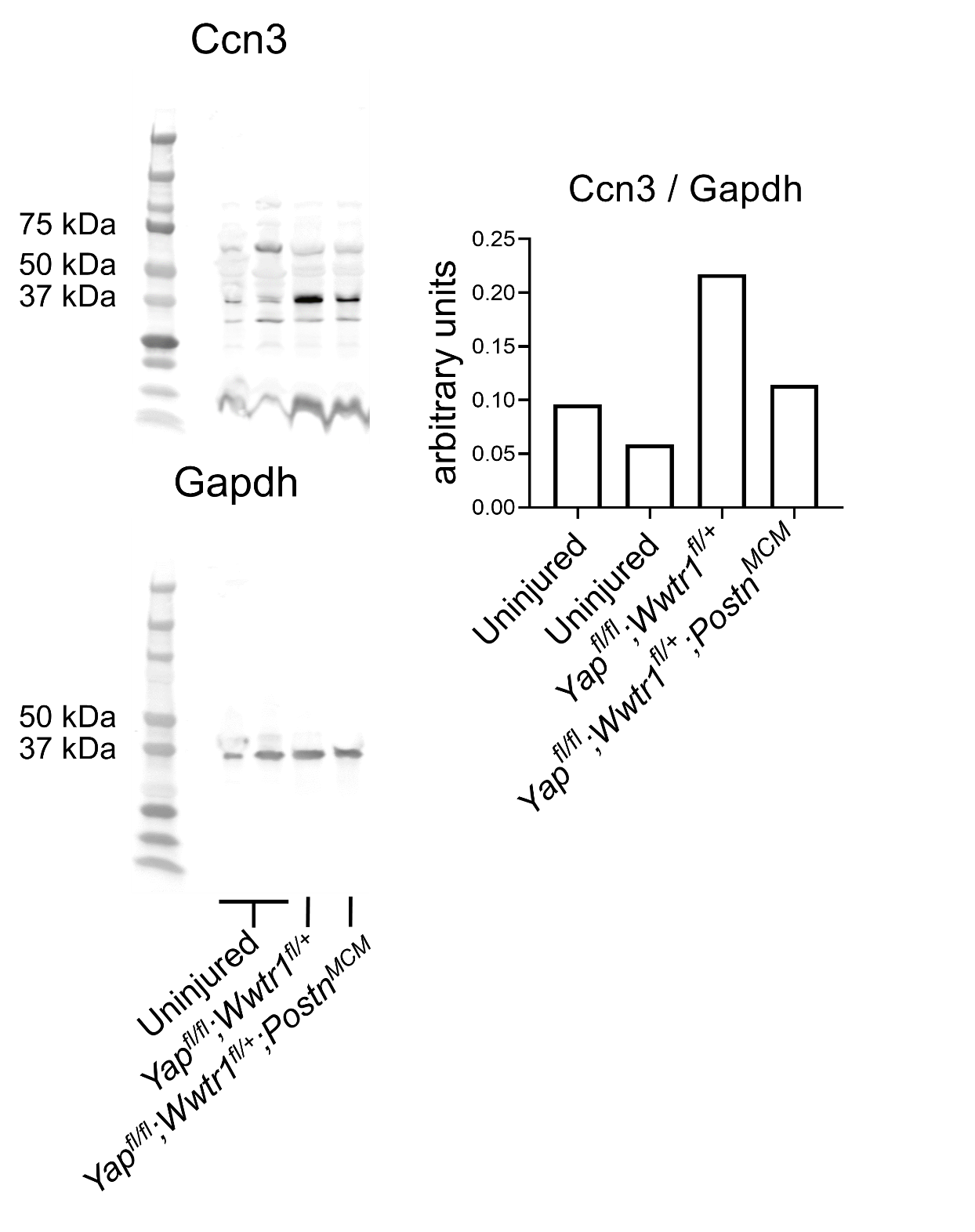

**Figure S12. Cardiac Ccn3 expression post injury.** Western blots illustrating expression of Ccn3 and Gapdh in left ventricles from uninjured and 14 dpi mice. Both Ccn3 and Gapdh expression are imaged on the same blot utilizing uniquely distinguishing secondary antibodies (IRDye® 800CW and IRDye® 680RD respectively), however due to similar masses separate images were required. Each lane represents expression from an individual mouse. Data presented in Figure 7C are derived from lanes 2, 3, and 4. Graph denotes densitometry of Ccn3 relative to Gapdh.

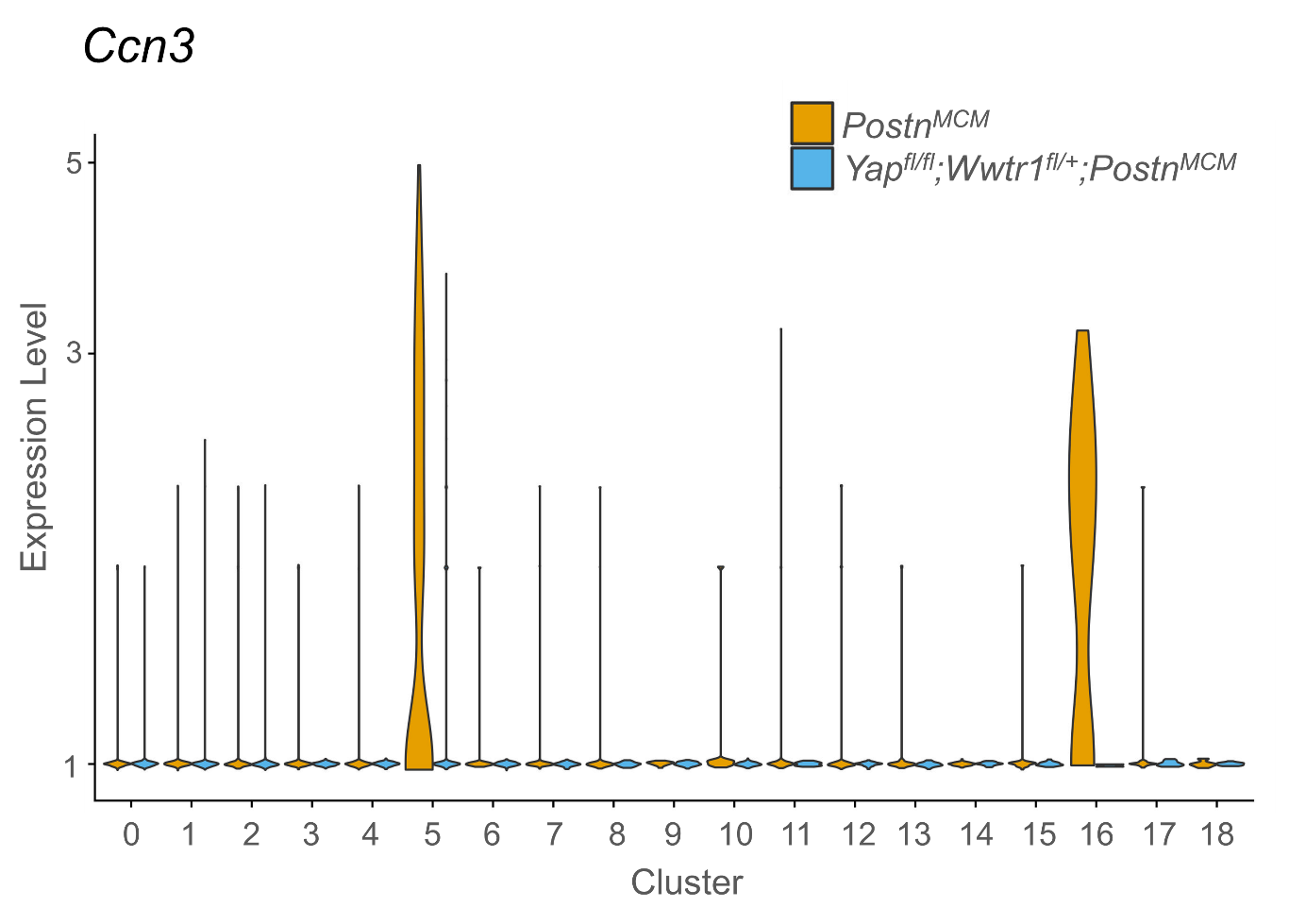

**Figure S13. CCN3 expression in cardiac fibroblasts is dependent on Yap/Wwtr1 expression.** Violin plots denoting UMI count for *Ccn3* for interstitial cell clusters, between *R26-eGFP^f/+^;Postn^MCM^* or *Yap^fl/fl^;Wwtr1^fl/+^;Postn^MCM^*  adult mice 7 dpi. Clusters are in reference to Figure 5A.

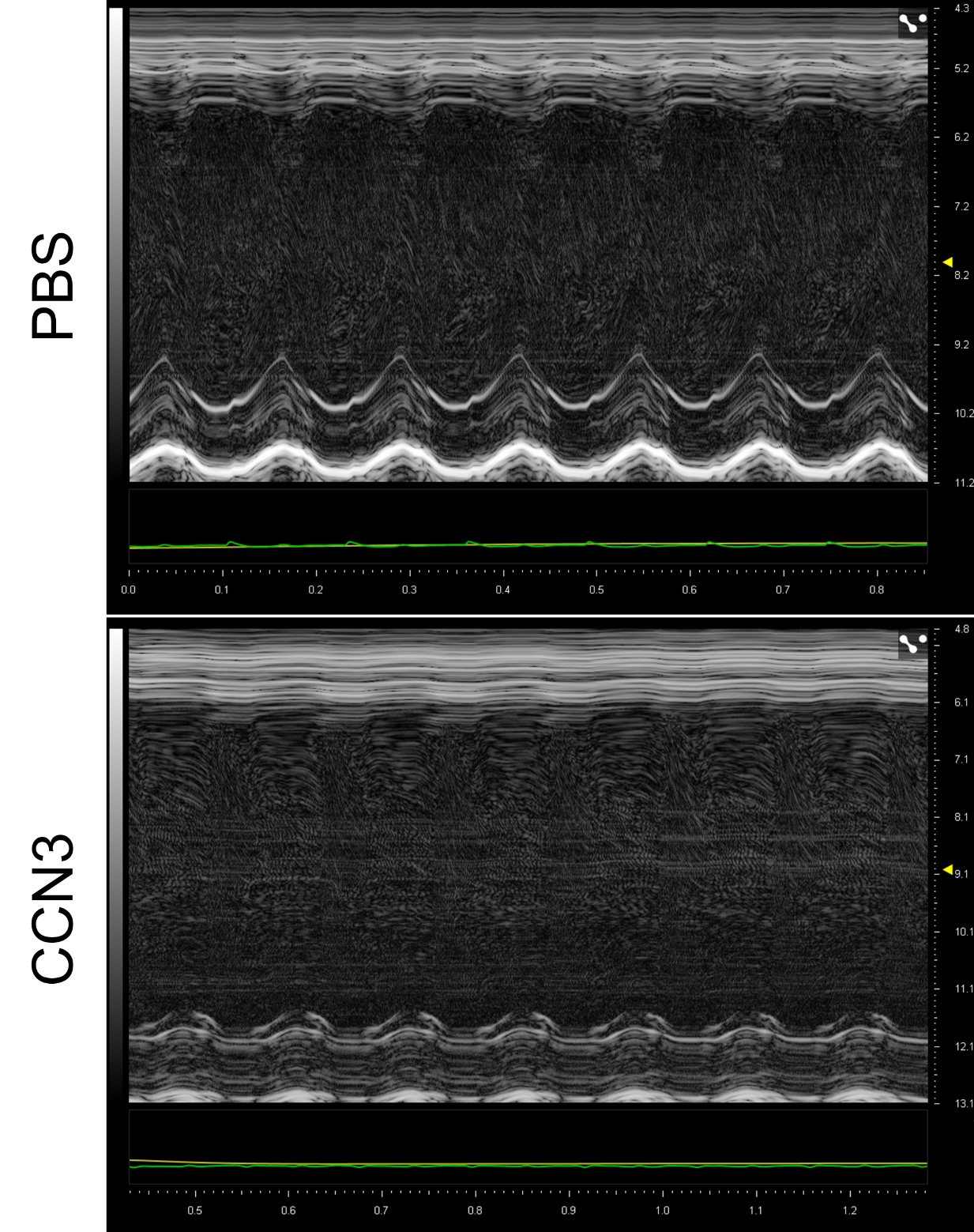

**Figure S14. CCN3 treatment reduces cardiac function after myocardial infarction.** Representative M-mode echocardiograms of left ventricles from PBS or CCN3 treated 28 dpi. Short axis view at mid papillary muscle. Horizontal scale indicates seconds of trace. Vertical scale indicates mm of depth from probe.

**Supplementary Tables**

**Table S1. Antibodies Used.**

| **Antibody** | **Use** | **Manufacturer** | **Concentration Used** |
| --- | --- | --- | --- |
| Anti-GFP [α-GFP] (ab13970) | IHC Primary | Abcam | 1:1000 |
| Anti-Yap [α-Yap] (#4912) | IHC Primary/ Western Blotting | Cell Signaling | 1:500 |
| Anti-Taz [α-Wwtr1] (560235) | IHC Primary/ Western Blotting | BD Pharmingen | 1:500 |
| IgY (H+L) Goat anti-Chicken, Alexa Fluor® 488 (A11039) | IHC Secondary | Invitrogen | 1:500 |
| Anti-Cardiac Troponin T antibody [α-Ctnnt] [1C11] (ab8295) | IHC Primary | Abcam | 1:500 |
| Goat anti-Rabbit IgG (H+L) Highly Cross-Adsorbed Secondary Antibody, Alexa Fluor 555 (A21429) | IHC Secondary | Invitrogen | 1:500 |
| Goat anti-Mouse IgG (H+L) Highly Cross-Adsorbed Secondary Antibody, Alexa Fluor Plus 555 (A32727) | IHC Secondary | Invitrogen | 1:1000 |
| 0IgG (H+L) Cross-Adsorbed Goat anti-Rabbit, Alexa Fluor® 488 (A11008) | IHC Secondary | Invitrogen | 1:1000 |
| Anti-CCN3 [α-Ccn3] antibody (ab137677) | Western Blotting | Abcam | 1:500 |
| Anti-Gapdh [α-Gapdh] (MAB5718) mouse monoclonal | Western Blotting | R&D Systems | 1:500 |
| Anti-Gapdh [α-Gapdh] (2118S) rabbit monoclonal | Western Blotting | Cell Signaling | 1:500 |
| IRDye® 800CW Goat anti-Mouse IgG (H + L) | Western Blotting | Licor | 1:20000 |
| IRDye® 800CW Goat anti-Rabbit IgG (H + L) | Western Blotting | Licor | 1:20000 |
| IRDye® 680RD Goat anti-Mouse IgG (H + L) | Western Blotting | Licor | 1:20000 |
| IRDye® 680RD Goat anti-Rabbit IgG (H + L) | Western Blotting | Licor | 1:20000 |

**Table S2. siRNA reagents.**

| **siRNA Reagent** | **Target** | **Source** |
| --- | --- | --- |
| MISSION® siRNA Universal Negative Control #1 - SIC001-10NMOL | Negative Control | Sigma |
| MISSION® siRNA Universal Negative Control #2 - SIC002-10NMOL | Negative Control | Sigma |
| SASI_Rn01_00114054 | *Yap* | Sigma |
| SASI_Rn02_00203876 | *Yap* | Sigma |
| SASI_Rn01_00120445 | *Wwtr1* | Sigma |
| SASI_Rn01_00120446 | *Wwtr1* | Sigma |

**Table S3. qRT-PCR primer sequences.**

| **Gene** | **Primer** | **Sequence** | **Organism** |
| --- | --- | --- | --- |
| *18s* | Forward | TAGTTGGATCTTGGGAGCGG | *Rattus norvegicus* |
| *18s* | Reverse | TAGAACCGCGGTCCTATTCC | *Rattus norvegicus* |
| *Yap* | Forward | TACATAAACCATAAGAACAAGACCACA | *Rattus norvegicus* |
| *Yap* | Reverse | GCTTCACTGGAGCACTCTGA | *Rattus norvegicus* |
| *Wwtr1* | Forward | CCCAATCTTGCGATGAATCACC | *Rattus norvegicus* |
| *Wwtr1* | Reverse | CTCCATCGGATCCTCTGAAGC | *Rattus norvegicus* |

**Table S4. Marker gene expression across UMAP clustering of interstitial cardiac cells at 7 days post injury.**

**Table S5. Differential gene expression within the cardiac fibroblasts subcluster of interstitial cells from myofibroblast Yap and Wwtr1 depleted and control hearts.** Differential genes identified by P < .05. Log2-fold change indicates relative expression in cardiac fibroblasts from *Yap^fl/fl^;Wwtr1^fl/+^;Postn^MCM^* hearts.

**Table S6. Marker gene expression across cardiac fibroblast UMAP subcluster at 7 days post injury.**
